# Influence of ploidy and genetic background on stress tolerance of intraspecific yeast hybrids

**DOI:** 10.1101/2025.11.08.686703

**Authors:** Kaisa Rinta-Harri, Tino Koponen, Dominik Mojzita, Paula Jouhten, Gianni Liti, Kristoffer Krogerus

**Affiliations:** VTT Technical Research Centre of Finland, Tekniikantie 21, 02150, Espoo, Finland; Aalto University, School of Chemical Engineering, Department of Bioproducts and Biosystems, Kemistintie 1, 02150 Espoo, Finland; IRCAN, CNRS, INSERM, Côte d’Azur University, Nice, France

**Keywords:** yeast, breeding, heterosis, ploidy, stress tolerance

## Abstract

Hybrid vigor, or heterosis, is widely exploited in yeast strain improvement. Yet, how ploidy and genetic background jointly shape heterosis across industrially relevant stresses remains unclear. Here, we generated 1023 *Saccharomyces cerevisiae* intraspecific hybrids derived from 18 genetically diverse parents, using two different approaches, yielding sets of hybrids with variable ploidy for the same parental combinations. High-throughput growth assays in media with five stress conditions (14% ethanol, 1.5 M NaCl, 0.15 M lactic acid, 0.05 M acetic acid, 0.05 M HMF) revealed extensive heterosis across 6138 hybrid-condition combinations. Most combinations displayed mid-parent heterosis and over a third exceeded the best parent, with the strongest gains during growth in the presence of 14% ethanol and 1.5 M NaCl. Increasing ploidy was generally associated with reduced growth and reduced best-parent heterosis, whereas greater predicted hybrid heterozygosity or genetic distance between parents was positively associated with heterosis in the presence of 14% ethanol and 1.5 M NaCl. Domestication status also affected these trends, as crosses between two domesticated strains tended to perform better in the presence of ethanol and NaCl, while crosses between two wild strains grew best in control conditions and in the presence of acetic acid. Together, these results demonstrate condition-dependent contributions of ploidy and parentage to heterosis and provide targeted breeding strategies for the improvement of stress-tolerance in industrial yeasts.

## Introduction

*Saccharomyces cerevisiae* is one of the most important industrial microbes, and it has a central role in food fermentation and production of biofuels, pharmaceuticals, enzymes and chemicals (Borodina and Nielsen 2014). Yeast strains are constantly developed to introduce novel features and to improve process and cost efficiency. Synthetic biology has greatly advanced the speed and decreased the cost of strain development. Still, there is a need for more robust host strains that synthetic biology can build on, and complex traits, such as stress tolerance, are challenging to improve through metabolic engineering. For example, lactic acid, the monomer of polylactic acid bioplastics, can be industrially produced from starch with yeast, but the use of complex second generation lignocellulosic raw materials is still a challenge, especially at low pH (Marchesan et al. 2021). Breeding has been exploited for strain development and in studies on yeast genetics for decades (Krogerus et al. 2017; Steensels et al. 2014b), with early work dating back to the 1930s (Winge and Laustsen 1938). Crossbred strains often show qualities superior to those of both parents, and this phenomenon is known as hybrid vigour or heterosis. Indeed, numerous recent studies have demonstrated that a wide range of industrially-relevant traits can be improved through yeast breeding (Krogerus et al. 2015; Mertens et al. 2015; Peris et al. 2017; Snoek et al. 2015; Steensels et al. 2014a). These include fermentation rates and yields, stress tolerance, and secondary metabolite production. This phenomenon is not unique to yeast and has been exploited to significantly improve animal and crop yields. Today, most of the world’s maize, sorghum and sunflower is produced from hybrid plants (Hochholdinger and Yu 2025).

While heterosis has been exploited for strain development through yeast breeding, the mechanisms that govern this important phenomenon are not fully understood. Factors that contribute to heterosis include: complementation of deleterious mutations through increased heterozygosity (Plech et al. 2014; Shapira et al. 2014), interactions between heterozygous alleles (Shapira and David 2016), and regulatory incompatibilities (Herbst et al. 2017; Tirosh et al. 2009). Yeast breeding is also coupled with a simultaneous increase in genome size (or ploidy). In *S. cerevisiae*, for example, haploid cells of opposite mating type can mate to form a diploid cell (Merlini et al. 2013). Ploidy, which influences gene dosage, heterozygosity, and cell size, also has an impact on yeast fitness, and may contribute to heterosis (Charron et al. 2019; Krogerus et al. 2016; Takagi et al. 1983; Zavaleta et al. 2024; Zörgö et al. 2013). However, the link between ploidy and heterosis has not been studied at large scale and rarely using polyploid parent strains and hybrids.

Recent population-scale genomic surveys of wild and domesticated *S. cerevisiae* strains, have revealed that polyploidy is enriched in specific human-associated populations (e.g. industrial brewing, distilling and baking strains), suggesting that polyploidy may provide a competitive advantage in certain stressful environments (Fischer et al. 2021; Gallone et al. 2016; Peter et al. 2018). Ploidy is also positively correlated with heterozygosity in both domesticated and wild *S. cerevisiae* strains (Fischer et al. 2021; Vittorelli et al. 2025). Heterozygosity can positively affect fitness through both dominance complementation and occurrence of superior (overdominant) heterozygous loci (Hallin et al. 2016; Plech et al. 2014; Shapira and David 2016; Shapira et al. 2014). Increased ploidy levels are also linked to increased genome instability, and this instability and plasticity allows polyploid strains to more rapidly adapt to stressful environments (Marsit et al. 2021; Selmecki et al. 2015).

Yeast breeding relies upon the ability of the yeast cells to form mating-competent cells with a single mating type (Merlini et al. 2013). Here, we exploit two different strategies of generating mating-competent cells to construct a large set of intraspecific yeast hybrids with variable ploidy between 18 genetically distinct parent strains. The first, more traditional, approach relies on conversion to heterothallism and sporulation. Here, the ploidy of the mating-competent cells is typically halved, and cells undergo meiotic recombination. Because of the reliance on sporulation, it is only applicable to fertile strains. Many industrial yeast strains, such as those in Ale beer, French dairy, and Sake clades (De Chiara et al. 2022), are sterile, and are hence often excluded from large-scale breeding studies. The second approach, in contrast, utilises CRISPR/Cas9 to force a mating-type change in the strain of interest (Krogerus et al. 2021a; Xie et al. 2018). This approach yields mating-competent strains with the original ploidy and genome intact. Furthermore, it can be applied to sterile strains as well. These two approaches were combined to effortlessly construct a set of 1023 distinct intraspecific hybrids for 153 pairwise crosses. Growth of the hybrid and parent strains in various high-stress media was compared to determine the extent of heterosis among the hybrids. The influence of ploidy and genetic distance between parent strains on growth and heterosis was also explored, with the aim of improving predictability in yeast breeding.

## Materials & Methods

### Yeast strains

Eighteen genetically diverse parent strains of *S. cerevisiae* (Table 1) were selected from previous studies (Krogerus et al., 2021; Peter et al., 2018). Auxotrophic, mating-competent variants of these strains were generated for hybrid creation using two strategies. In the first, strains were directly converted to stable *MATa* or *MATalpha* strains using CRISPR/Cas9 plasmids containing protospacer sequences targeting either *MATalpha* or *MATa*, respectively (Krogerus et al. 2021a). In the second approach, strains were first made heterothallic by deleting the endonuclease-coding *HO* gene using CRISPR/Cas9, after which stable *MATa* or *MATalpha* strains were obtained following sporulation and ascospore dissection. *ho* deletion strains were transferred to 1% potassium acetate agar for sporulation. After 7 days of incubation at 25 °C, ascospores were digested using Zymolyase 100 T (US Biological, Salem, MA, USA) and dissected on YPD agar using the MSM400 dissection microscope (Singer Instruments, Roadwater, UK). Mating-type was confirmed with PCR as described below. The stable *MATa* and *MATalpha* strains from both approaches were converted into lysine and uracil auxotrophs, respectively, by deleting *LYS2* and *URA3* using CRISPR/Cas9. Auxotrophy was confirmed by testing their inability to grow in minimal medium (0.67% yeast nitrogen base with ammonium sulphate, 1% glucose). All mating-competent strains are listed in Supplementary Table S1.

**Table 1.**
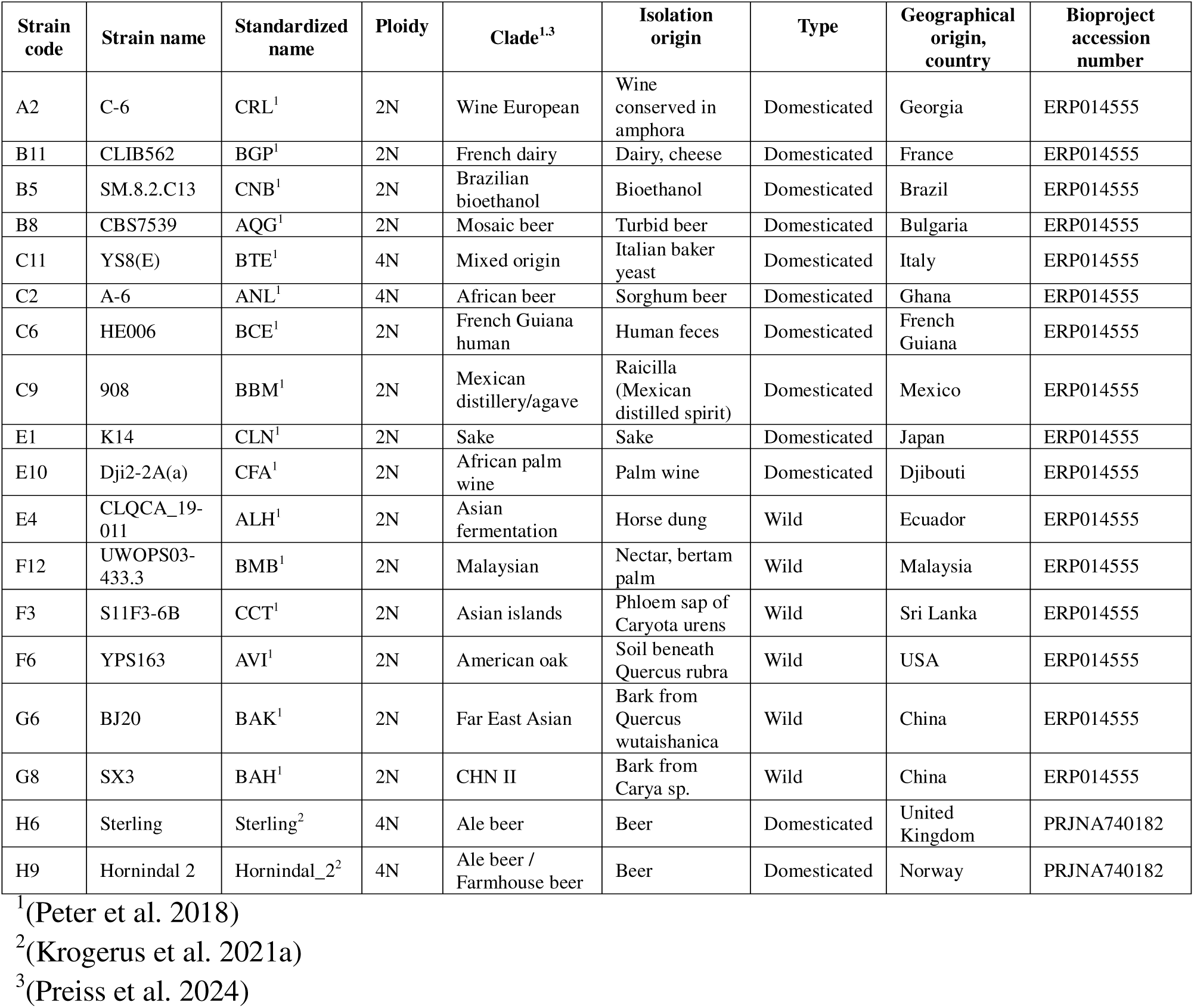
Summary of parent strains.

For creating intraspecific hybrids, the 64 auxotrophic mating-competent strains (31 *MATa* and 33 *MATalpha*) were grown overnight in 150 µL Yeast Peptone Dextrose (YPD) medium in round-bottom 96-microwell plates (Thermo Scientific, Denmark) at 26 °C. A total of 1023 pairwise crosses between the *MATa* and *MATalpha* strains were carried out as follows: 1 µL aliquots of each parent strain were transferred adjacently and mixed on YNB agar (0.67% yeast nitrogen base with ammonium sulphate, 1% glucose) in a 4×4 grid (16 crosses per plate). The agar plates were incubated at 26 °C for up to five days. Prototrophic colonies emerging on the agar plate were assumed to be successful hybrids and were streaked onto fresh YNB agar. Successful hybridization was also confirmed by PCR as described below. All hybrid strains are listed in Supplementary Table S2.

### Cas9 plasmid construction and yeast transformations

Cas9 plasmids for mating-type switching and *HO* deletion, containing yeast codon-optimized Cas9 expressed under *TDH3p*, single-guide RNA (sgRNA) expressed under *SNR52p*, and *hygR* marker for selection on hygromycin were constructed previously (Krogerus et al. 2021a). The gRNA protospacer sequences, GTTCTAAAAATGCCCGTGCT, CAAATCATACAGAAACACAG, and AAACATGTAAGGCTTCATTA, target *MAT***a**, *MATalpha*, and *HO,* respectively. Cas9 plasmids targeting *URA3* and *LYS2* were constructed by cloning sgRNAs into a multicopy, 2-micron-based, base plasmid containing the NAT marker for selection on nourseothricin, the Cas9 expression cassette, and modified sgRNA expression cassette. The base Cas9 plasmid was linearized inside the modified sgRNA expression cassette, after the *SNR52* promoter and before the tracrRNA coding region, and used for Gibson assembly (NEBuilder® HiFi DNA Assembly; NEB) with specific ssDNA fragments (Supplementary Table S3) to introduce target-specific crRNA sequences into the sgRNA gene. The assembled plasmids were transformed into *Escherichia coli* TOP10 by electroporation, and the plasmid correctness was confirmed by Sanger sequencing.

Yeast transformations were performed using an optimized stationary phase transformation protocol (Tripp et al. 2013). Overnight yeast cultures were pelleted and incubated with 100 mM DTT (dithiothreitol) for 20 min at 42 °C. A lithium acetate-based transformation mix was added, together with 1 μg of purified Cas9 plasmid and 10 μg of donor DNA/repair fragment (only for deletions; listed in Supplementary Table S3), and cells were transformed at 42 °C for 40 min. The transformed cells were selected on plates containing either 400 mg/L hygromycin B (Sigma-Aldrich, Darmstadt, Germany) or 100 mg/L nourseothricin (Jena Bioscience AB-102 XL, Jena, Germany). Successful mating-type changes or gene deletions were determined by PCR as described below. Colonies from selection plates were streaked for single colonies three successive times on YPD agar plates before further use to encourage plasmid loss, after which they were stored at -80 °C.

### PCR to confirm mating-type change, hybridizations, and gene deletions

Prior to PCR, DNA was extracted by suspending freshly grown colonies of the strains from agar plates in 10 mM NaOH and heating to 95 °C for 15 minutes. Following brief centrifugation, the resulting crude DNA extract was used directly as template in PCR reactions. All PCR primers are available in Supplementary Table S3. The mating-type locus was amplified via PCR utilizing previously established primers (Huxley et al. 1990). These primers amplify a 404 bp fragment for the *MATalpha* locus and a 544 bp fragment for the *MATa* locus. PCR reactions were conducted using Phusion Plus PCR Master Mix with HF Buffer (Thermo Scientific, Finland) with a primer concentration of 0.5 μM in 96 well semi-skirted PCR plates (4titude, Wotton, Surrey, UK). The PCR products were subsequently separated and visualized using an Agilent ZAG DNA Analyzer capillary electrophoresis system (Agilent, Santa Clara, CA, USA).

### High-throughput growth assays

Yeast strains were characterized using high-throughput 96-well plate cultivations. Pre-cultivations were first carried out in minimal media supplemented with uracil and lysine (0.67% yeast nitrogen base with ammonium sulphate, 20 mg/L uracil, 60 mg/L lysine, 1% glucose). Pre-cultures were carried out in 96-well plates with round-bottom wells containing 140 µL of media. Wells were inoculated with 10 µL of yeast from glycerol stocks, and the plates were incubated at 26 °C without shaking for 2 days.

Five different high-stress growth media, along with a control media, were prepared. The control medium was minimal medium supplemented with uracil and lysine (0.67% yeast nitrogen base with ammonium sulphate, 20 mg/L uracil, 60 mg/L lysine, 1% glucose). The five high-stress media were further supplemented with either 14% ethanol, 1.5 M sodium chloride, 0.15 M lactic acid, 0.05 M acetic acid, or 0.05 M hydroxymethylfurfural (HMF). To initiate the cultivations, 140 µL of each of the six media were transferred to three replicate round-bottom 96-well plates and inoculated with 5 µL of mixed yeast pre-culture. The plates were then incubated at 25 °C with 1000 rpm shaking and monitored using Biomek i7 liquid handler station (Beckman Coulter Life Science, Brea, CA, USA), coupled with a Cytomat 2C 470 LIN Shaker incubator (Thermo Scientific, Waltham, MA, USA) and a SpectraMax iD5 Multimode Microplate reader (Molecular Devices, San Jose, CA, USA). Absorbance at 600 nm (OD600) was measured in three-hour intervals over a cultivation period of 96 hours.

### Genome sequencing and analysis

Genomes of the 18 wild-type parent strains had been sequenced previously (Krogerus et al. 2021a; Peter et al. 2018). DNA was extracted from the mating-competent strains generated here using a Mag-Bind® Blood & Tissue DNA HDQ 96 Kit (M6399; Omega Bio-tek, Norcross, GA, USA). Short-read library preparation and sequencing of strains was performed at NovoGene (Munich, Germany). Sequencing was performed on an Illumina NovaSeq X Plus sequencer in a multiplexed shared-flow-cell run, producing 2 × 150 bp paired-end reads. Short reads were trimmed and filtered with fastp using default settings (version 0.20.1) (Chen et al. 2018). Trimmed reads were aligned to a *S. cerevisiae* S288C reference genome (Engel et al. 2014) using BWA-MEM (Li and Durbin 2009), and alignments were sorted and duplicates were marked with sambamba (version 0.7.1) (Tarasov et al. 2015). Variants were jointly called in all strains using FreeBayes (version 1.32) (Garrison and Marth 2012). Variant calling used the following settings: --min-base-quality 30 --min-mapping-quality 30 --min-alternate-fraction 0.25 – min-repeat-entropy 0.5 --use-best-n-alleles 70 -p 2. The resulting VCF file (Variant Call Format) was filtered to remove variants with a quality score less than 1000 and with a sequencing depth below 10 per sample using BCFtools (Li 2011). The genetic distance between pairs of parent strains was calculated using PLINK (version 1.9) (Purcell et al. 2007), run with the ‘--genome full’ flag, as the sum of IBS0 and IBS1.

For phylogenetic analysis, the variants were filtered to retain only single nucleotide polymorphisms and remove sites with a minor allele frequency less than 5%. The filtered single nucleotide polymorphism (SNP) matrix was converted to PHYLIP format (https://github.com/edgardomortiz/vcf2phylip). A random allele was selected for heterozygous sites. A maximum likelihood phylogenetic tree was generated using IQ-TREE (version 2.0.3; Nguyen et al. 2015) run with the ‘GTR+G4’ model and 1000 bootstrap replicates (Minh et al. 2013).

### DNA content analysis by flow cytometry

Ploidy of the yeast strains was measured using SYTOX Green staining and flow cytometry (Haase and Reed 2002). Pre-cultures were started in 96-well plates from glycerol stocks by inoculating 140 µL of minimal medium with 5 µL from each of stock. Plates were incubated overnight at 26 °C without shaking. 10 µL of cell suspension from each well was transferred to a fresh 96-well plate containing 100 µL of ice-cold 77% (v/v) ethanol. The cells were fixed at -20 °C and stored in the freezer for up to 3 weeks until later analysis. Following fixation, cells were pelleted by centrifugation at 3000×*g* for 10 minutes, and the supernatant was carefully removed. The pellet was washed with 100 µL of 50 mM sodium citrate buffer (pH 7.2). The pellet was then resuspended in 66 µL of 50 mM sodium citrate buffer containing 0.25 mg/mL RNase A (Thermo Scientific, Vilnius, Lithuania) and 1 mg/mL proteinase K (Thermo Scientific, Vilnius, Lithuania). After overnight incubation at 37 °C, 33 µL of 50 mM sodium citrate buffer containing 6 µM SYTOX Green (Invitrogen, Eugene, OR, USA) was added. The cells were incubated overnight in the dark at 4 °C to allow for optimal staining. Flow cytometry analysis was conducted at the HiLife Flow Cytometry Unit at the University of Helsinki, using a NovoCyte Quanteon Flow Cytometer (Agilent, San Diego, CA, USA). 10000 events were collected from each sample, and fluorescence was monitored using the FITC channel, with an excitation wavelength of 488 nm and emission detection wavelength of 525 nm. The wild-type parent strains with known ploidy served as controls, and a standard curve was generated based on the fluorescence intensity of the control strains.

### Data processing

The absorbance data collected from the high-throughput cultivations was fitted to the ‘Gompertz’ function using the ‘QurveE’ package in R (Wirth et al. 2023). This function provided estimates of key growth parameters, including maximum specific growth rate (µ). Area under curve (AUC) was calculated from the fitted growth curves using the ‘gcplyr’ package in R (Blazanin 2024). Following curve fitting, averages of the three replicate measurements were calculated for the growth parameters. The median coefficient of variations between replicates across the whole data set were 5.5% and 6.9% for AUC and µ, respectively.

Heterosis of the hybrid strains was calculated based on AUC and µ. Mid-parent heterosis (MPH), meaning the hybrid performs better than the average of the parent strains, was calculated as follows:

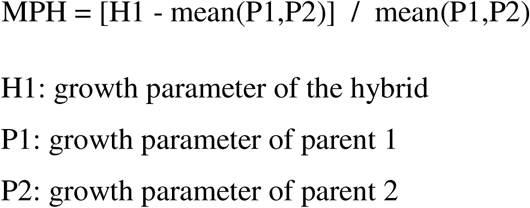

Best-parent heterosis (BPH), meaning the hybrid performs better than the best-performing parent, was calculated as follows:

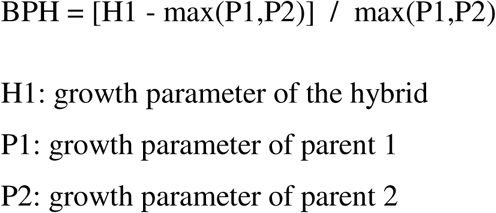

All data was plotted in R using ‘ggpubr’ and ‘pheatmap’ packages (Kassambara 2025; Kolde 2015). Linear regression models predicting heterosis (AUC.BPH) as a function of genetic distance between parent strains (G) and measured ploidy (P) were fit with the ‘lm’ function in R (AUC.BPH ∼ G + P + 0), while standardised β coefficients were calculated from the fit models with the ‘lm.beta’ package. Statistical analysis was performed using ‘rstatix’ and ‘agricolae’ packages in R (De Mendiburu 2017; Kassambara 2023). *p*-values from multiple comparisons were corrected with the Benjamini-Hochberg procedure. All growth data is available in Supplementary Table S4.

## Results

### Generating a large collection of intraspecific hybrids with variable ploidy

Eighteen genetically and geographically diverse *S. cerevisiae* strains were selected from collections of previously sequenced strains to be used as parent strains in the breeding trials (Krogerus et al. 2021a; O’Donnell et al. 2023; Peter et al. 2018). The eighteen strains were selected to represent the major clades described in Peter et al. (2018). The strains were selected at random for each clade, however, strains with publicly available long-read assemblies and known to form viable spores were preferred (De Chiara et al. 2022; O’Donnell et al. 2023). Out of the eighteen selected strains, twelve were domesticated and six wild strains, while four strains were tetraploid and the rest diploid (Table 1).

After the set of parent strains had been selected, auxotrophic variants with single mating types and variable ploidy were generated using two approaches (Figure 1A-B). The first, utilized CRISPR/Cas9-aided mating type switching to convert the mating type of the parents from *MATa*/*MATalpha* to either *MATa* or *MATalpha* (Krogerus et al. 2021a; Xie et al. 2018). This approach should, in theory, maintain the original ploidy and full genome of the parent strain. In the second approach, the *HO* gene was deleted from the parent strains to prevent spontaneous mating type switching after sporulation. The *ho* deletion strains were then sporulated, and viable spore clones were screened for a single mating type using PCR. This approach should, in theory, halve the ploidy and result in loss of heterozygosity of the parent strain. A total of 64 mating-competent variants were ultimately obtained (31 *MATa* and 33 *MATalpha* strains), with three of the original 18 parent strains not forming viable spores. The *MATa* and *MATalpha* strains were further converted into lysine and uracil auxotrophs, respectively, through deletion of *LYS2* and *URA3* with CRISPR/Cas9. This was done to enable direct selection of successful hybrids.

**Figure 1.**
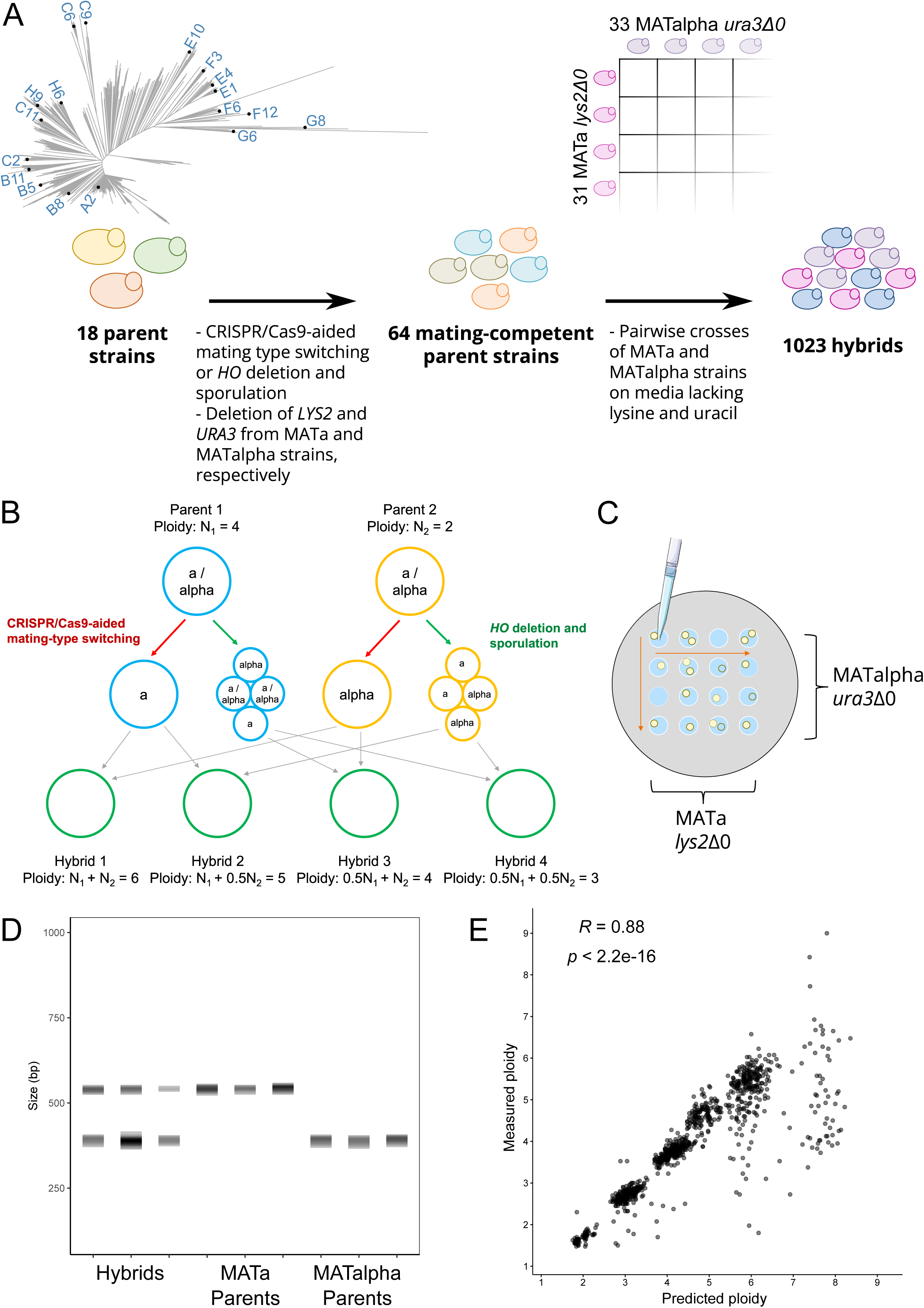
An overview of the method used to generate intraspecific hybrids in this study. (**A**) A set of 64 mating-competent variants were derived from 18 genetically diverse parent strains. The neighbour-joining phylogenetic tree shows the genetic relationship between the parent strains in comparison to the 1,011 yeast genomes strains (Peter et al. 2018). Pairwise crosses of the mating-competent strains were performed to obtain a set of 1023 distinct hybrids. (**B**) A schematic over how the method yields hybrids of different ploidy from crosses of the same parent strains. (**C**) Hybridization was carried out by mixing droplets of cell suspension from two different mating-competent strains directly on agar plates with media lacking lysine and uracil. (**D**) Successful hybridization was confirmed using mating-type PCR. Parents produced a single band for either *MATa* or *MATalpha*, while hybrids produced both bands. PCR products were separated using capillary electrophoresis. (**E**) Correlation between predicted and measured ploidy values in the hybrid strains. Predicted ploidy is calculated as the sum of the parent strain ploidy. *R* is the Pearson correlation coefficient.

The ploidy of the mating-competent strains was determined using SYTOX Green staining and flow cytometry, and for the majority of the strains, it corresponded to what was expected based on the ploidy of the wild-type parent strains (Supplementary Table S1). When CRISPR/Cas9-aided mating type switching was used, only the ploidy of those derived from wild-type strain F12 were altered (doubling from diploid to tetraploid for both the *MATa* and *MATalpha* variants). When *ho* deletion and sporulation was used, the ploidy was expectedly halved from that of the wild-type strains in most of the strains, with exceptions for some variants from wild-type strains C6, F3, G6, H6, and H9. Whole-genome sequencing of all 64 mating-competent strains was also performed to ensure that no major changes had occured in the genomes compared to the 18 original wild-type strains. The number of single nucleotide polymorphisms identified in the mating-competent strains compared to the wild-type strain from which they were derived ranged from 0 to 41, with an average of 10 (Supplementary Table S1).

Once the set of 64 mating-competent strains was assembled, pairwise crosses between all the *MATa* and *MATalpha* strains were carried out (Figure 1A-C). Crosses were performed on solid growth media lacking uracil and lysine. As the mating-competent strains were auxotrophic, any prototrophic colonies emerging on the agar plates were considered successful hybrids between the two mating-competent parent strains. To further confirm that hybridization had been successful, the mating-type of the hybrids was confirmed to be *MATa*/*MATalpha* using PCR (Figure 1D). All 1023 attempted crosses (33 × 31) yielded successful hybrids. As the two strategies for generating mating-competent strains yielded strains of variable ploidy derived from the same parent strain, hybrids of variable ploidy were also obtained for each of the crosses between the original 18 parent strains (Figure 1B). For example, crosses between mating-competent strains derived from a tetraploid and diploid parent were expected to yield a series of hybrids with ploidy 3N, 4N, 5N and 6N. In general, the measured ploidy of the hybrids corresponded well (*r* = 0.88) with the predicted ploidy, i.e., the sum of the ploidies of the parent strains (Figure 1E). Interestingly, at higher predicted ploidy (≥ 6), the measured ploidy was often lower than expected, highlighting the genomic instability at high ploidy levels. The mean ploidy of the hybrid strains was 3.9.

### Genetic background and ploidy influences growth of intraspecific hybrids

Microtiter cultivations were performed with the 18 wild-type strains, 64 mating-competent strains, and 1023 hybrid strains in various stress media. The base growth media consisted of YNB with 1% glucose, which was then supplemented with either 0.05 M acetic acid, 0.05 M 5-hydroxymethylfurfural (HMF), 0.15 M lactic acid, 1.5 M NaCl, or 14% ethanol. Growth was monitored by regular turbidity measurements at 600 nm, and the maximum specific growth rate (µ) and the area under curve (AUC) were determined for each strain in each media by fitting to the Gompertz growth model. Variation in growth (both µ and AUC) in the different stress media was observed among the wild-type strains (Supplementary Figure S1). In general, the growth of the mating-competent strains correlated strongly (*r* ranging from 0.59 to 0.94) to that of the wild-type strains they were derived from in all media (Supplementary Figure S2). However, in the stress-supplemented media, the growth of the mating-competent strains was typically lower than that of the wild-type strains. This impairment was likely caused by their auxotrophy (Swinnen et al. 2015). Variation in growth in the different stress media was also observed among the hybrid strains depending on which parent strain they were derived from (Figure 2A). Out of the five stress media, growth was weakest in the media with 1.5 M NaCl and 0.15 M lactic acid. In all growth media, there was a positive correlation (*r* ranging from 0.52 to 0.83) between the observed growth (as AUC) in the parent strains and the hybrids grouped by parent strain (Figure 2B), indicating, expectedly, that the performance of the hybrids is linked to the performance of the parent strains. Similar positive correlations were observed based on the maximum specific growth rate (Supplementary Figure S3).

**Figure 2.**
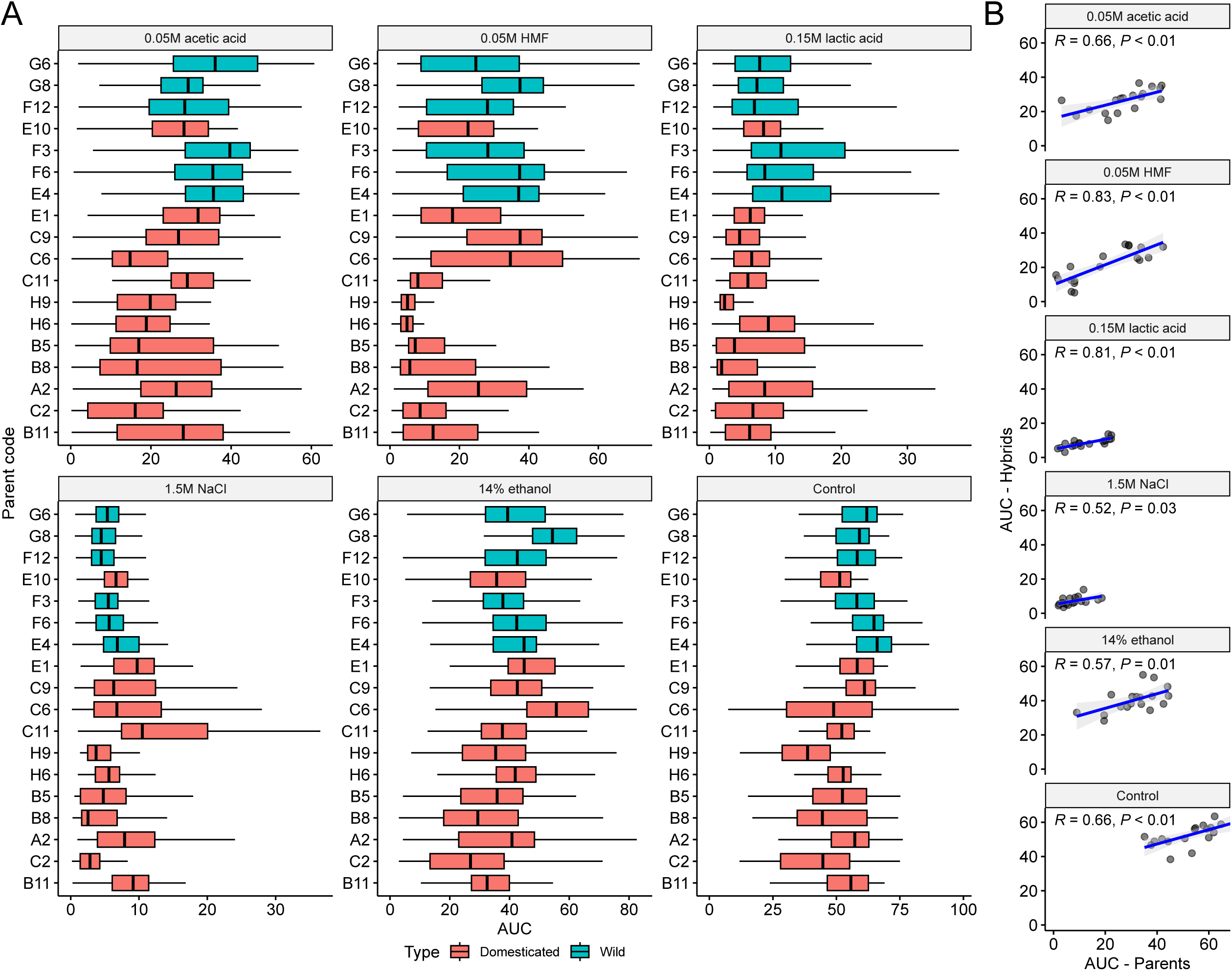
Growth of the hybrid strains in different media. (**A**) Boxplots of the area under curve (AUC) in the hybrids as grouped by parent strain in the six different growth media. Strains are ordered based on the phylogenetic relationship between the parents (maximum likelihood phylogenetic tree based on SNPs at 105677 sites, rooted with strain G8 as outgroup). Boxes are colored cyan if the parent strain is considered wild, red if considered domesticated. (**B**) The correlation between AUC in the parent strains and hybrids grouped by parent strain in the different growth media. *R* is the Pearson correlation coefficient.

Overall, when comparing growth of the hybrids with that of the parent strains, no significant difference in the mean maximum specific growth rate or AUC was observed in the control media or the media supplemented with acetic acid, HMF or NaCl (Figure 3A and Supplementary Figure S4A). However, in the medium supplemented with ethanol, both significantly higher maximum specific growth rate and AUC were observed for the hybrids. In contrast, significantly higher maximum specific growth rate and AUC were observed for the parent strains in the medium supplemented with lactic acid. This suggested that heterosis was more common in ethanol stress than in lactic acid stress. Within the set of hybrid strains, significant differences in mean AUC and maximum specific growth rate could also be seen when hybrids were grouped based on whether the parent strains were domesticated or wild (Figure 3B and Supplementary Figure S4B). In all growth media, except that supplemented with NaCl, hybrids between two domesticated parent strains performed worse than any having at least one wild parent. Hybrids between two wild strains tended to have the highest growth. In the media supplemented with NaCl, however, an opposite trend was observed, with a slightly higher AUC for hybrids between two domesticated strains. This compared well with what was observed with the parent strains, where domesticated strains showed the highest growth in this medium (Supplementary Figure S1C). No significant differences in maximum specific growth rate were observed though in the medium supplemented with NaCl (Supplementary Figure S4B).

**Figure 3.**
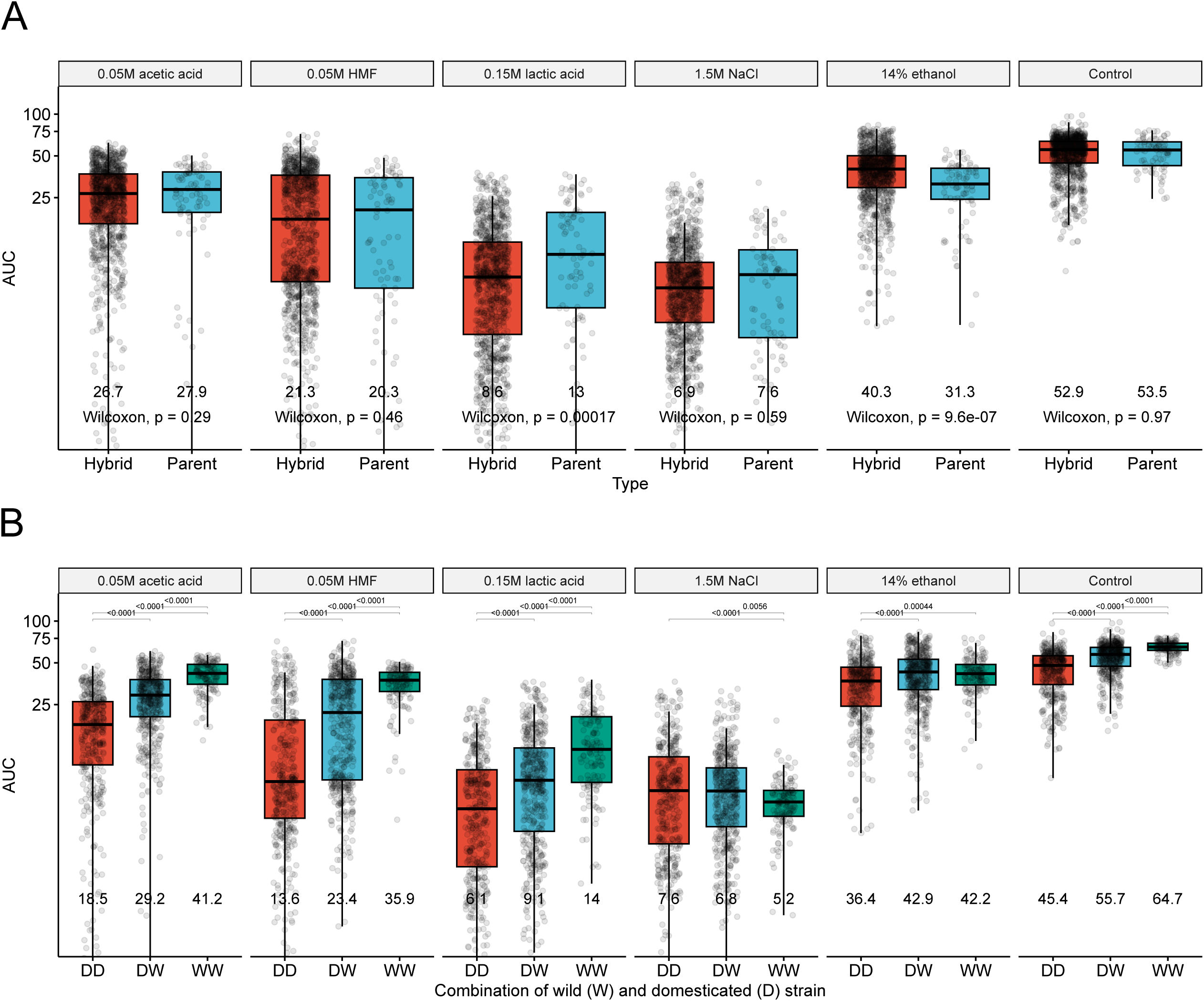
Growth comparison of the hybrid and parent strains in different media. (**A**) Boxplots of the area under curve (AUC) in the hybrid (red) and parent (blue) strains in the six different growth media. Mean values of each group are displayed under the boxes. Statistical difference between the hybrid and parent strains in each media was tested by an unpaired Wilcoxon test. (**B**) Boxplots of the area under curve (AUC) in the hybrid strains as grouped by parent strain combination in the six different growth media. DD (red): both parent strains were domesticated, DW (blue): one parent strain was domesticated, and one was wild, WW (green): both parent strains were wild. Statistical difference between the groups in each media was tested by one-way ANOVA and Tukey’s post-hoc test, and *p*-values are shown above the boxes.

Hybridization is typically associated with an increase in ploidy, and here we generated sets of variable ploidy hybrids from the same parent strain crosses to investigate how ploidy influences growth in various environments. Growth of the hybrids showed a moderate negative correlation (*r* ranging from -0.22 to -0.57) with increasing ploidy in all growth media (Figure 4A and Supplementary Figure 5A), indicating that a higher ploidy is generally detrimental to growth. Growth of hybrids strains with high ploidy (>4.5) was particularly poor in the medium supplemented with 0.05 M HMF. However, this group of hybrids was also overrepresented (56.4% of the bin) by those created from either of the parent strains H6 and H9, which also performed poorly in this medium (Figure 2A). When comparing the relationship between ploidy and growth of the hybrids in the different media based on the parent strains, a distinction between hybrids made from domesticated and wild strains could be seen (Figure 4B and Supplementary Figure 5B). Hybrids made from wild strains tended to have a stronger negative correlation between ploidy and growth. Interestingly, a positive correlation between ploidy and growth in multiple growth media could even be seen among the hybrids derived from parent strain H9, a tetraploid farmhouse brewing strain.

**Figure 4.**
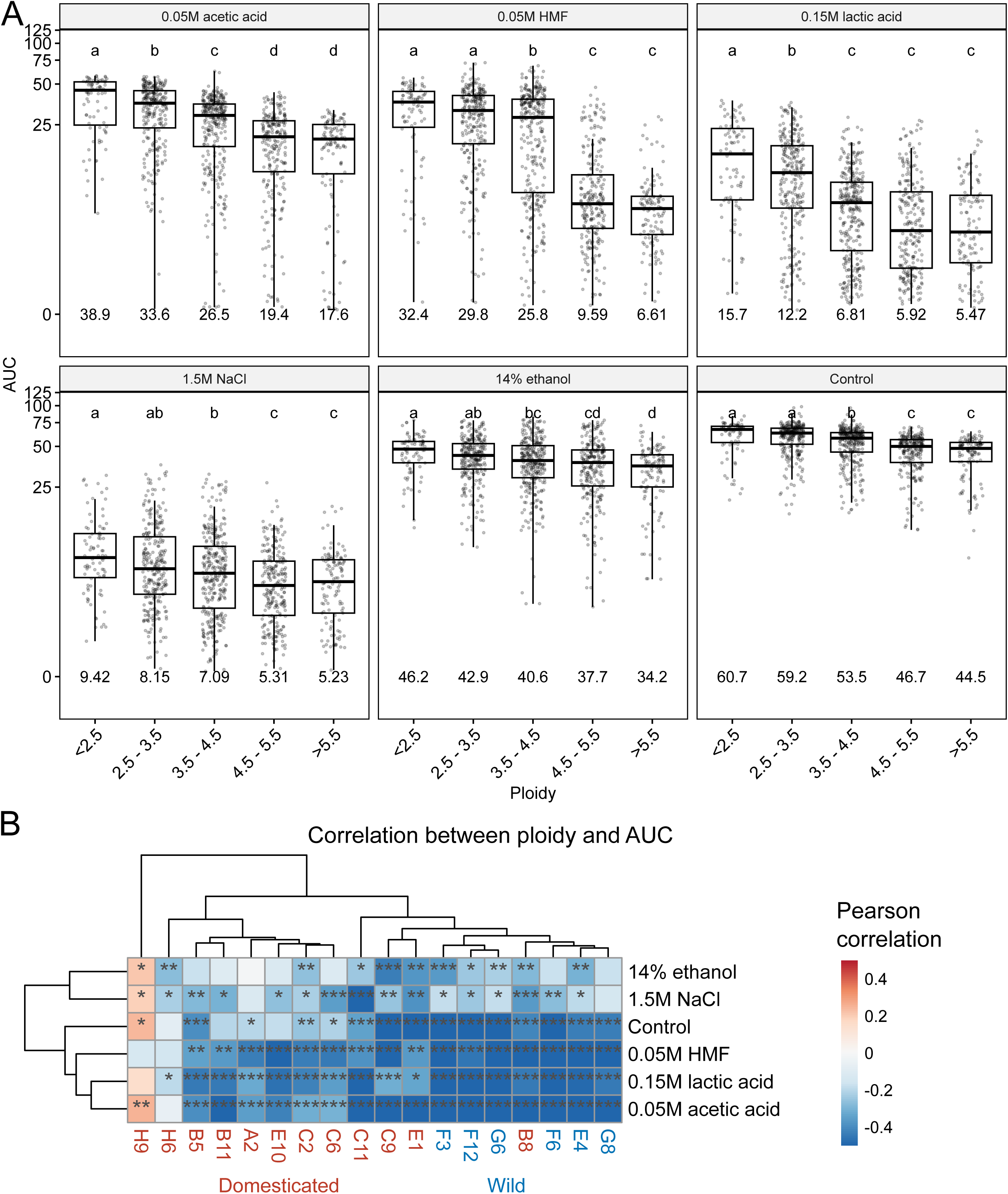
Influence of ploidy on the growth of the hybrids in different media. (**A**) Boxplots of the area under curve (AUC) in the hybrid strains as grouped by ploidy in the six different growth media. The mean value of each group is shown below the boxes. Different letters above the boxes within each growth media indicate significant differences (*p* < 0.05) as determined by one-way ANOVA and Tukey’s post-hoc test. (**B**) A heatmap visualizing the Pearson correlation coefficient between ploidy and AUC in the hybrids as grouped by parent strain and growth media. Red and blue colors indicate a positive and negative correlation coefficient, respectively. The *p*-values from multiple comparisons were corrected using the Benjamini-Hochberg procedure. Asterisks indicate the *p-*values as follows: * *p* < 0.05, ** *p* < 0.01, and *** *p* < 0.001.

### The impact of ploidy and genetic distance between parent strains on heterosis is dependent on growth conditions

While no significant differences in the average growth between the parent and hybrid strains were observed in most media, hybrid strains often outperformed the parent strains from which they were derived. Out of the 6138 cultivations that were carried out with 1023 hybrids in six different media, positive best-parent heterosis was observed in 2140 cases (34.9%) and positive mid-parent heterosis in 3302 cases (53.8%) when calculated based on AUC. Values were similar (2064 and 3166, respectively) when calculated based on the maximum specific growth rate. The frequency of heterosis was dependent on the growth media, with positive best-parent heterosis based on AUC being most prevalent in the medium supplemented with 14% ethanol and least prevalent in that with 0.15 M lactic acid (Figure 5A). These were also the two media where the largest relative difference in mean growth was observed between the parent and hybrid strains (Figure 3A). The prevalence of positive best-parent heterosis was also dependent on parent strain. Among the hybrid strains showing positive best-parent heterosis, those created from parent strains C9, F3, and C6 were overrepresented and present in a frequency higher than expected based on their frequency among all hybrid strains (Figure 5B). Parent strains C9 and C6 were both isolated in Latin America, belonging to the sister clades ‘Mexican agave’ and ‘French Guiana human’. Heterosis was rarer, on the other hand, among hybrids created from parent strains A2, B5, C2, and H9, which include strains used for wine and beer fermentations.

**Figure 5.**
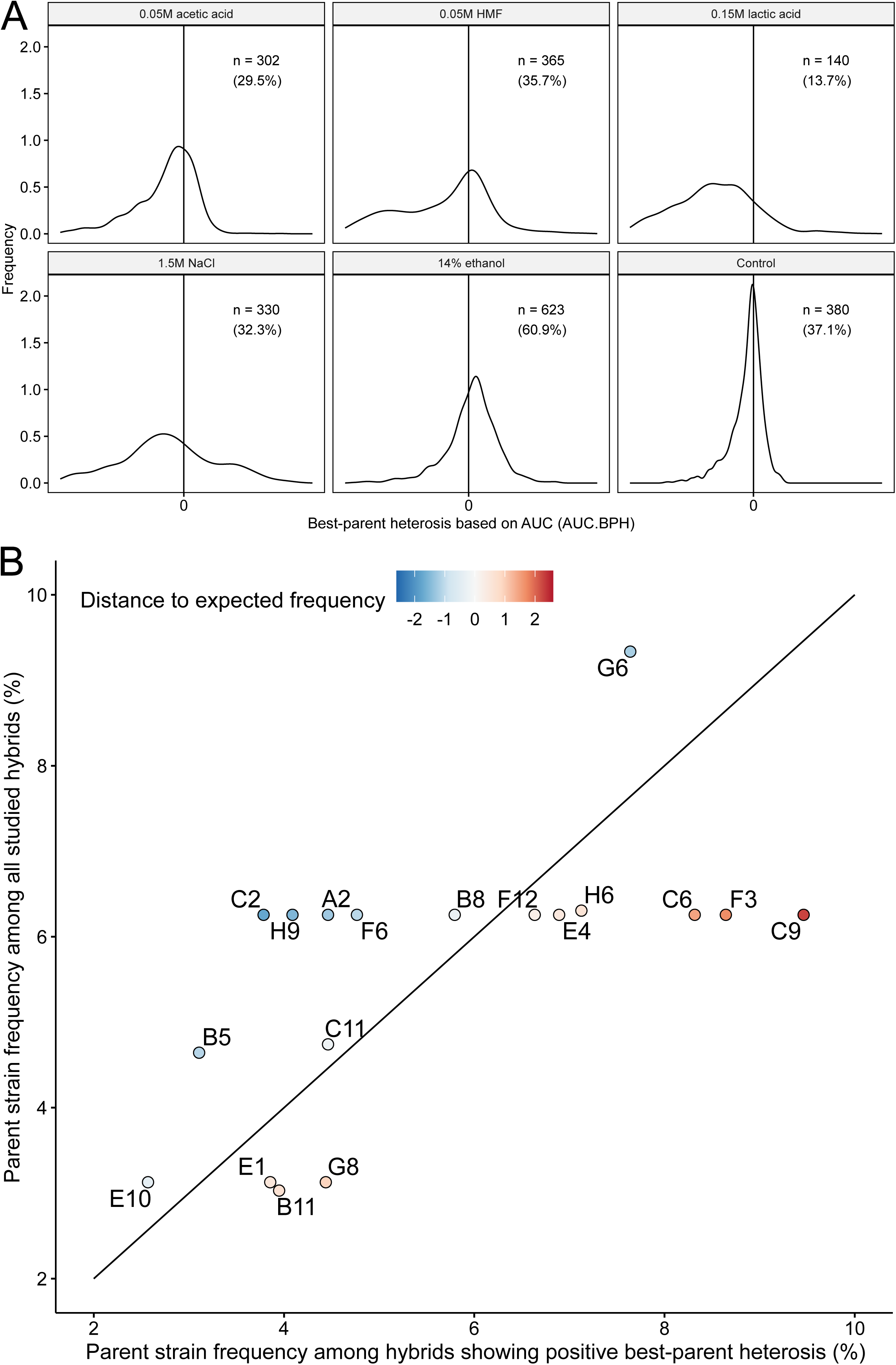
Frequency of best-parent heterosis in the hybrid strains. (**A**) Density plots of best-parent heterosis values based on area under curve (AUC.BPH) in the hybrid strains in the different growth media. The vertical line is at AUC.BPH = 0. The n-value indicates the number of strains (and percentage of all hybrid strains) where AUC.BPH was larger than 0, i.e. hybrids outperformed both parent strains. (**B**) A comparison of the parent strain frequency among the hybrids showing positive best-parent heterosis and all hybrid strains. The diagonal line indicates values where the frequencies are equal. Parent strains to the right of the diagonal are overrepresented among the hybrids with best-parent heterosis, while strains to the left are underrepresented. Points are colored based on the distance to the expected frequency (i.e. diagonal line).

As with growth, best-parent heterosis tended to correlate negatively with an increasing ploidy in the hybrids (Figure 6A). However, the correlations were weaker than the ones observed between AUC and ploidy (*r* here ranging from 0.014 to -0.27). In the media containing 1.5 M NaCl or 14% ethanol, no significant differences in best-parent heterosis were observed between the different ploidy bins. On the other hand, in the control medium, and those supplemented with 0.05 M acetic acid or HMF, best-parent heterosis decreased at increasing hybrid ploidy. Varying effects of increasing ploidy on best-parent heterosis were also observed when hybrids were grouped based on parent strain (Figure 6B). Again, as observed above for the correlation to growth, a distinction between hybrids made from domesticated and wild strains could be seen, as hybrids made from wild strains tended to have a stronger negative correlation between ploidy and best-parent heterosis. Significant positive correlation between ploidy and heterosis in different growth media could be seen among the hybrids made from parent strains H9 and B11, a tetraploid farmhouse brewing and diploid French dairy strain, respectively. Furthermore, weak positive correlation between ploidy and heterosis could also be observed in other strains derived from beverage fermentation, such as H6 (ale beer), B8 (mosaic beer), and E10 (African palm wine).

**Figure 6.**
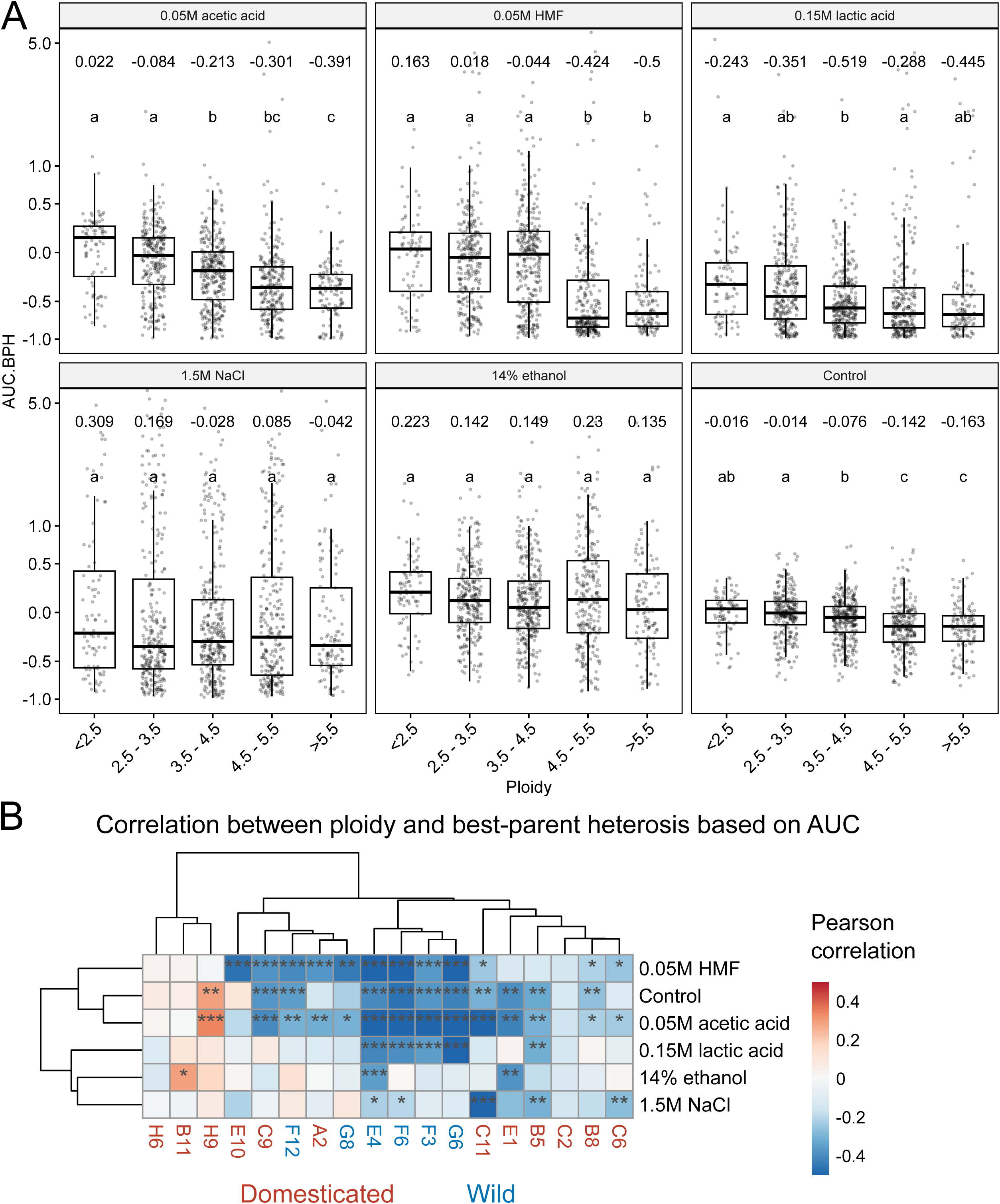
Influence of ploidy on the best-parent heterosis based on area under curve (AUC.BPH) in different media. (**A**) Boxplots of the AUC.BPH in the hybrid strains as grouped by ploidy in the six different growth media. The mean value of each group is shown above the boxes. Different letters above the boxes within each growth media indicate significant differences (*p* < 0.05) as determined by one-way ANOVA and Tukey’s post-hoc test. (**B**) A heatmap visualizing the Pearson correlation coefficient between ploidy and AUC.BPH in the hybrids as grouped by parent strain and growth media. Red and blue colors indicate a positive and negative correlation coefficient, respectively. The *p*-values from multiple comparisons were corrected using the Benjamini-Hochberg procedure. Asterisks indicate the *p-*values as follows: * *p* < 0.05, ** *p* < 0.01, and *** *p* < 0.001.

In addition to an increased genome size, hybridization is also typically associated with an increased heterozygosity. While whole-genome sequencing of the hybrids was not carried out here, heterozygosity in the hybrids and genetic distance between the parent strains was estimated based on sequencing data from the mating-competent parent strains. Increased best-parent heterosis was observed in the media supplemented with 1.5 M NaCl or 14% ethanol for hybrids with an increasing amount of predicted heterozygous sites, either from high initial heterozygosity in either of the parents or from high genetic distance between the parent strains (Figure 7A). In the other media, an increased predicted heterozygosity in the hybrids appeared to have no effect on heterosis. As the predicted number of heterozygous sites correlates very strongly with the genetic distance between the parents (Supplementary Figure S6), very similar correlation coefficients were expectedly obtained for best-parent heterosis and genetic distance between parents (Figure 7B). Hence, if the goal is to increase tolerance towards osmotic and ethanol stress, then exploiting hybridization between genetically distant parent strains appears to be effective. When best-parent heterosis was visualized as a function of both genetic distance between parent strains and ploidy (Figure 8A), the detrimental impact of increasing ploidy on heterosis could be seen in many media (e.g. 0.05M acetic acid, 0.05M HMF, and 0.15M lactic acid), while a positive impact of increasing genetic distance between parent strains could not be as easily observed. Indeed, in simple linear regression models predicting heterosis as a function of these two variables, ploidy tended to have a larger impact on heterosis than genetic distance (larger magnitude of β coefficient) in almost all media (Figure 8B). Only in the media supplemented with 14% ethanol, did genetic distance have a larger impact on predicted heterosis than ploidy, as also indicated by the correlation between best-parent heterosis and genetic distance between parent strains in this media. However, prediction of heterosis based on these two variables alone was poor (*R^2^* values ranging from 0.06 to 0.48).

**Figure 7.**
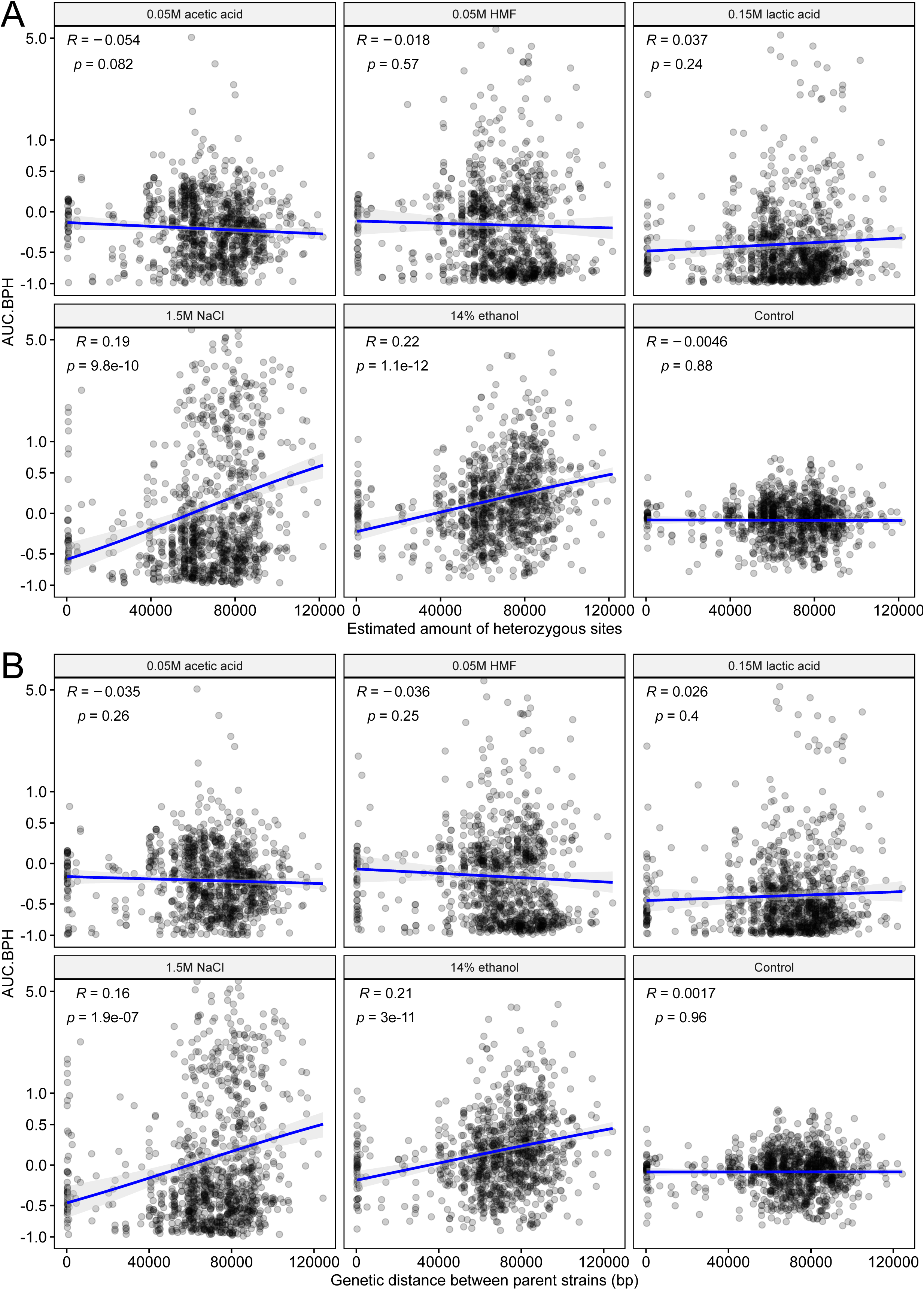
Influence of heterozygosity and genetic distance on the best-parent heterosis based on area under curve (AUC.BPH) in different media. (**A**) The correlation between the estimated amount of heterozygous sites in the hybrids and AUC.BPH. Heterozygous sites were estimated based on the single nucleotide polymorphisms (SNPs) detected in the mating-competent parent strains of each hybrid. (**B**) The correlation between the genetic distance between the parent strains and AUC.BPH. Genetic distance was estimated based on the sum of IBS0 and IBS1 alleles in each pairwise cross as estimated in *PLINK* based on SNPs detected in the mating-competent parent strains of each hybrid. (**A-B**) *R* is the Pearson correlation coefficient.

**Figure 8.**
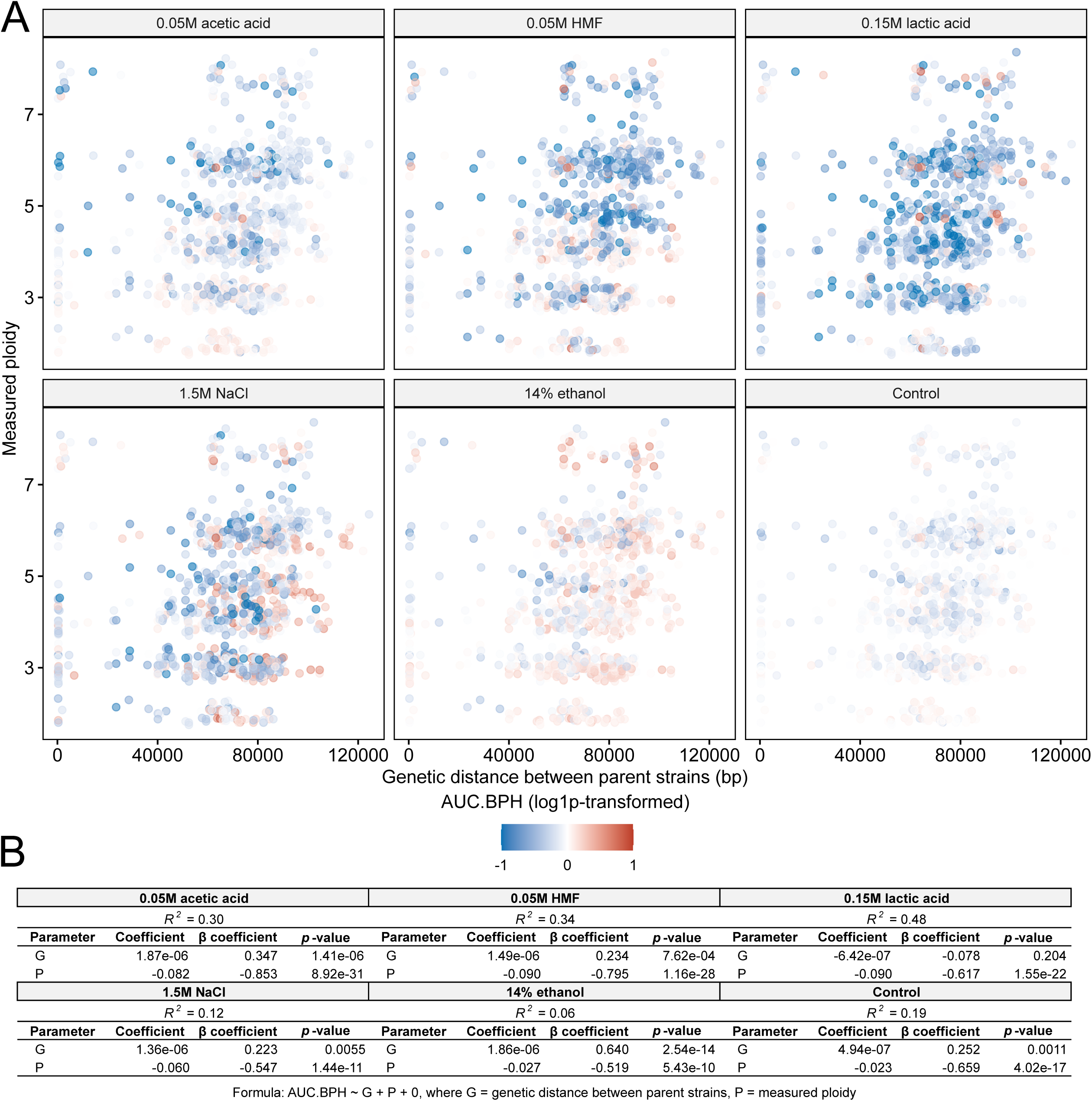
– Combined influence of genetic distance and ploidy on the best-parent heterosis based on area under curve (AUC.BPH) in different media. (**A**) Scatter plots of the genetic distance between the parent strains and measured ploidy in the hybrids. Points are colored based on the AUC.BPH value (log1p-transformed; AUC.BPH values remain positive or negative even after transformation). Genetic distance was estimated based on the sum of IBS0 and IBS1 alleles in each pairwise cross as estimated in *PLINK* based on SNPs detected in the mating-competent parent strains of each hybrid. (**B**) Regression coefficients, standardized β coefficients, and associated *p*-values for the linear regression models predicting log1p-transformed AUC.BPH as a function of genetic distance between parent strains (G) and measured ploidy (P) (AUC.BPH ∼ G + P + 0).

The effect of genetic distance on heterosis also varied depending on parent strain (Figure 9A and Supplementary Figure S7). As was similarly seen between ploidy and heterosis, there was a weak positive correlation between heterosis and genetic distance between parent strains when hybrids were made with nearly all the domesticated parent strains across all growth media. In hybrids made with wild parent strains, on the other hand, there was a significant negative correlation to heterosis with genetic distance in multiple media and parent strains. Heterosis was also impacted by the parent strain combination (Figure 9B and Supplementary Figure S7). The highest average best-parent heterosis values in the media containing 1.5M NaCl and 14% ethanol were observed for hybrids made between two domesticated strains. Conversely, in the control media and that supplemented with 0.05M acetic acid, hybrids between two wild strains had higher average best-parent heterosis values than hybrids between two domesticated strains.

**Figure 9.**
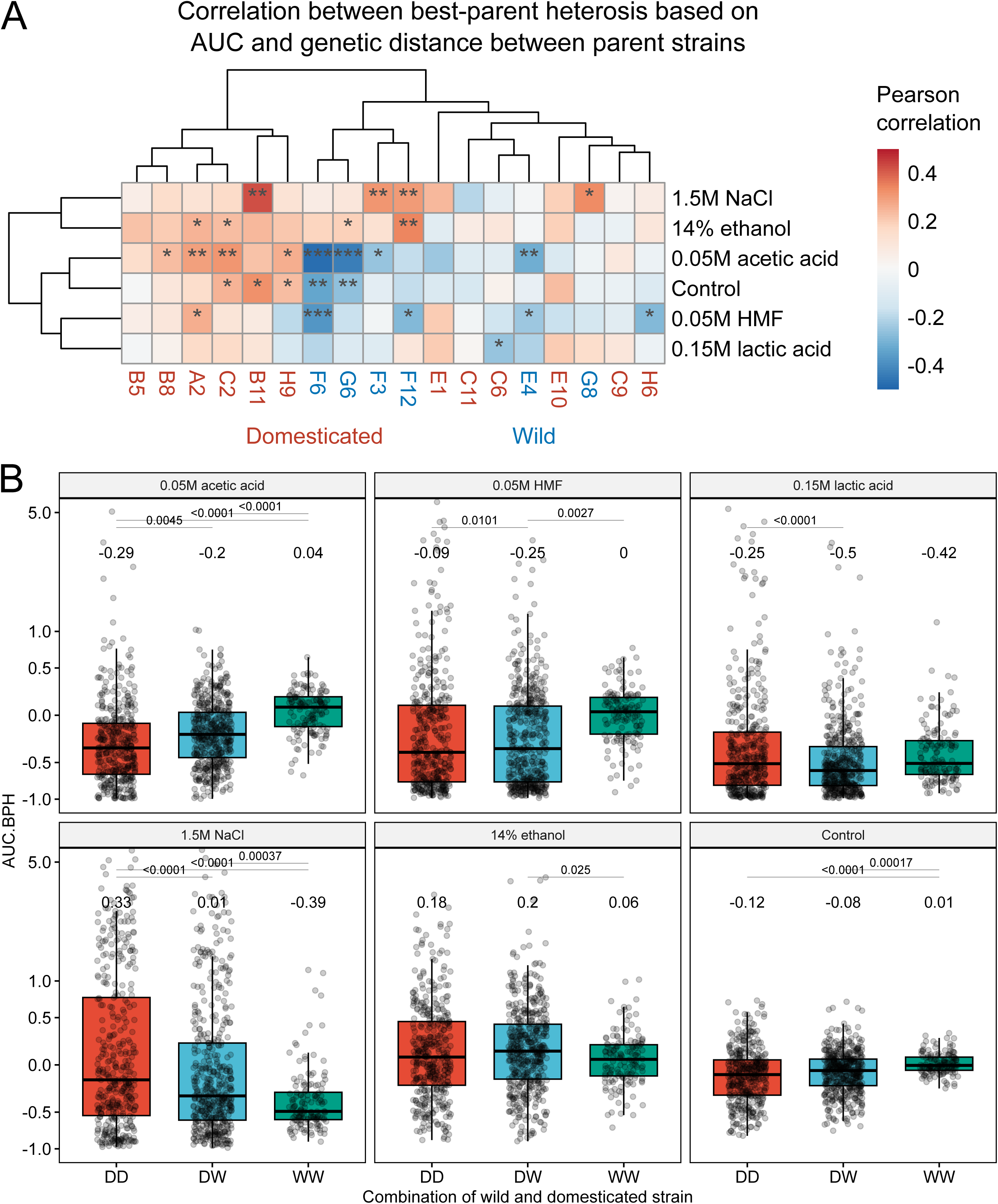
Influence of genetic distance and parent strain on the best-parent heterosis based on area under curve (AUC.BPH) in different media. (**A**) A heatmap visualizing the Pearson correlation coefficient between genetic distance between parent strains and AUC.BPH in the hybrids as grouped by parent strain and growth media. Red and blue colors indicate a positive and negative correlation coefficient, respectively. The *p*-values from multiple comparisons were corrected using the Benjamini-Hochberg procedure. Asterisks indicate the *p-*values as follows: * *p* < 0.05, ** *p* < 0.01, and *** *p* < 0.001. (**B**) Boxplots of the AUC.BPH in the hybrid strains as grouped by parent strain combination in the six different growth media. DD (red): both parent strains were domesticated, DW (blue): one parent strain was domesticated, and one was wild, WW (green): both parent strains were wild. The mean value of each group is shown above the boxes. Statistical difference between the groups in each media was tested by one-way ANOVA and Tukey’s post-hoc test, and *p*-values are shown above the boxes.

## Discussion

Creating large sets of hybrids has traditionally been challenging and time-consuming, particularly in cases where parent strains are sterile. In this study, we demonstrate how such a set of 1000+ intraspecific hybrids can be rapidly constructed from a library of stable mating-competent strains. By using two different approaches to create the mating-competent strains, series’ of hybrids with variable ploidy could be obtained for each pairwise cross of parent strains. In general, the ploidy of the generated hybrids correlated well with what was predicted based on the ploidy of the parent strains. However, as has been observed in previous studies as well (Barker et al. 2025; Hirota et al. 2024; Peris et al. 2020), the genomes of the hybrids predicted to have high ploidy levels (>6N) appeared to be unstable. Hirota et al. showed that constructed polyploid yeast strains of up to 32N tended to quickly converge to ploidy below 4N (Hirota et al. 2024). Indeed, ploidy levels over tetraploid are very rare among the over 3000 natural *S. cerevisiae* isolates that have been tested (Loegler et al. 2024; Peter et al. 2018). While this instability is impractical for immediate applied use, it can potentially be exploited to further improve the strains through directed laboratory evolution. This approach was used to enhance the growth of an approximately allooctaploid interspecies hybrid in glucose- and xylose-based complex media (Peris et al. 2020).

Hybrids are typically exploited for their potential to exhibit heterosis. Here, we observed best-parent heterosis in approximately 35% of the cultivations that were carried out with the hybrids. This was in line with previous studies, where similar frequencies were reported (Plech et al. 2014; Shapira et al. 2014). The effect of ploidy on heterosis has, to our knowledge, not been studied extensively in yeast before. Here, growth in all environments, and best-parent heterosis in most environments, negatively correlated with increasing ploidy. A similar negative impact of increasing genome size on growth was recently reported in interspecific yeast hybrids (Peris et al. 2020). In plants, triploid and tetraploid hybrids often show higher heterosis than diploid hybrids, but as observed here, growth rate tends to decrease at higher ploidy (Fort et al. 2016; Hallahan 2024; Miller et al. 2012). Hybridization typically also results in increased heterozygosity, which in plants is an important contributor to heterosis. In yeast, studies have shown that there is minimal correlation between genetic distance between parents and heterosis (Bernardes et al. 2017; Plech et al. 2014; Shapira et al. 2014). Plech et al. (2014) did, however, observe that sequence divergence between parents and heterosis did correlate in domesticated strains. In most of the tested growth conditions, we also observed no correlation, but there was a weak positive correlation between increased genetic distance between the parent strains and heterosis during growth in ethanol and osmotic stress. It must be mentioned, that here, whole-genome sequencing was only performed on the mating-competent parent strains, not on the hybrids. Hence, only predictions on the number of heterozygous sites in the hybrids could be made. Particularly in the high ploidy hybrids, where measured ploidy was lower than predicted, it is likely that significant loss of heterozygosity had occurred. Furthermore, we only generated and characterized intraspecific hybrids, but it is likely that heterosis and the relationship with genetic distance would be more prevalent in interspecific hybrids as previously demonstrated (Bernardes et al. 2017). While not tested, CRISPR/Cas9-aided mating-type switching should be applicable to other *Saccharomyces* yeasts provided new protospacer sequences are designed.

In applied fermentations, studies that examine the effect of ploidy in hybrids on fermentation performance have been limited. In work with interspecific *Saccharomyces* hybrids, using an *S. cerevisiae* parent strain from the ‘Mosaic beer’ clade, improved fermentation rate in brewer’s wort was achieved using allotetraploid hybrids, compared to triploid or diploid ones (Krogerus et al. 2016; Krogerus et al. 2021b; Nikulin et al. 2018). In more recent work, using a more genetically diverse set of *S. cerevisiae* parents, no significant correlation was observed between ploidy and fermentation performance in beer (Zavaleta et al. 2024). Here, we saw a weak correlation between heterosis and increasing ploidy levels in hybrids that were made from tetraploid brewing strains, such as H6 and H9. Improved resistance to hydrolysate inhibitors has also been demonstrated in a triploid strain compared to an isogenic diploid one (Liu et al. 2021). An increase in ploidy results in an increased cell size up to a certain limit (Barker et al. 2025; Fukuda 2023). Yeast cells with high ploidy levels and larger cell size are more viable at increased osmolarity and ethanol concentrations (Barker et al. 2025; Dinh et al. 2008), which could explain why heterosis here was common even at high ploidy in the 1.5 M NaCl and 14% ethanol media.

With the developments in metabolic engineering and synthetic biology, it has become possible to introduce novel complex pathways to yeast. However, most engineering studies are performed in host strains from a limited pool of laboratory strains. These strains are not representative of the phenotypic diversity available in yeast and might not be suitable for industrial application compared to domesticated strains (Gallone et al. 2016; Peter et al. 2018). Recent studies have shown that superior yields from heterologous pathways can be obtained by using non-laboratory strains (Long et al. 2024; Zhu et al. 2025). Zhu et al. (2025) demonstrated that out of the five diverse host strains they tested, the highest limonene yields could be obtained from an engineered baking strain. Yeast breeding, especially if more predictable, could be a useful tool to obtain more stress-tolerant host strains. Here, hybrids showing best-parent heterosis were identified for all the stress-supplemented media.

To conclude, we demonstrate how large sets of intraspecific hybrids with variable ploidy can be easily generated, even from sterile strains. By characterizing these hybrids in different growth conditions, we demonstrate condition-dependent contributions of ploidy and parentage to heterosis and provide targeted breeding strategies for improvement of stress-tolerance in industrial yeasts. Here, a fairly limited number of stresses were tested, but the screening could readily be expanded to other growth conditions. From an applied perspective, tolerance to extreme temperatures is also of interest, and testing both heat and cold tolerance would be valuable. To improve predictability of heterosis and allow rational design of hybrid strains for targeted improvement of specific phenotypes, machine learning could be further applied to elucidate relationships between the strain background or genotypes and heterosis. Together, this study highlights the power of yeast breeding as a strain development tool and increases our understanding of how breeding can be used for targeted enhancement of stress tolerance.

## Declarations

## Acknowledgements

The authors thank Noora Aarnio (Biomedicum Flow Cytometry Unit, University of Helsinki) for her kind assistance during flow cytometry analysis. Mari Nyyssönen (VTT) for her kind assistance in extraction of DNA for whole-genome sequencing. Ronja Valajärvi (VTT) for technical assistance.

## Ethical approval

This article does not contain any studies with human participants or animals performed by any of the authors.

## Availability of data and material

Illumina sequencing reads have been deposited in NCBI-SRA under BioProject number PRJNA1344768 (https://www.ncbi.nlm.nih.gov/bioproject/).

## Authors’ contributions

KRH: Designed experiments, performed experiments, edited the manuscript

TK: Performed experiments, edited the manuscript

DM: Performed experiments, edited the manuscript

PJ: Conceived the study, edited the manuscript

GL: Conceived the study, edited the manuscript

KK: Conceived the study, designed experiments, performed experiments, performed data analysis, wrote the manuscript.

All authors read and approved the final manuscript.

## References

Barker J, Murray A, Bell SP (2025) Cell integrity limits ploidy in budding yeast. G3 (Bethesda) 15(2) doi:10.1093/g3journal/jkae286

Bernardes JP, Stelkens RB, Greig D (2017) Heterosis in hybrids within and between yeast species. J Evol Biol 30(3):538–548 doi:10.1111/jeb.13023

Blazanin M (2024) gcplyr: an R package for microbial growth curve data analysis. BMC Bioinformatics 25(1):232 doi:10.1186/s12859-024-05817-3

Borodina I, Nielsen J (2014) Advances in metabolic engineering of yeast Saccharomyces cerevisiae for production of chemicals. Biotechnology Journal 9(5):609–620 doi:10.1002/biot.201300445

Charron G, Marsit S, Hénault M, Martin H, Landry CR (2019) Spontaneous whole-genome duplication restores fertility in interspecific hybrids. Nature Communications 10(1):4126–4126 doi:10.1038/s41467-019-12041-8

Chen S, Zhou Y, Chen Y, Gu J (2018) fastp: an ultra-fast all-in-one FASTQ preprocessor. Bioinformatics 34(17):i884–i890 doi:10.1093/bioinformatics/bty560

De Chiara M, Barre BP, Persson K, Irizar A, Vischioni C, Khaiwal S, Stenberg S, Amadi OC, Zun G, Dobersek K, Taccioli C, Schacherer J, Petrovic U, Warringer J, Liti G (2022) Domestication reprogrammed the budding yeast life cycle. Nat Ecol Evol 6(4):448–460 doi:10.1038/s41559-022-01671-9

De Mendiburu F (2017) Package ‘agricolae’ Title Statistical Procedures for Agricultural Research. Statistical procedures for agricultural research

Dinh TN, Nagahisa K, Hirasawa T, Furusawa C, Shimizu H (2008) Adaptation of Saccharomyces cerevisiae cells to high ethanol concentration and changes in fatty acid composition of membrane and cell size. PLoS One 3(7):e2623 doi:10.1371/journal.pone.0002623

Engel SR, Dietrich FS, Fisk DG, Binkley G, Balakrishnan R, Costanzo MC, Dwight SS, Hitz BC, Karra K, Nash RS, Weng S, Wong ED, Lloyd P, Skrzypek MS, Miyasato SR, Simison M, Cherry JM (2014) The reference genome sequence of Saccharomyces cerevisiae: then and now. G3 (Bethesda, Md) 4(3):389–98 doi:10.1534/g3.113.008995

Fischer G, Liti G, Llorente B (2021) The budding yeast life cycle: More complex than anticipated? Yeast 38(1):5–11 doi:10.1002/yea.3533

Fort A, Ryder P, McKeown PC, Wijnen C, Aarts MG, Sulpice R, Spillane C (2016) Disaggregating polyploidy, parental genome dosage and hybridity contributions to heterosis in Arabidopsis thaliana. New Phytol 209(2):590–9 doi:10.1111/nph.13650

Fukuda N (2023) Apparent diameter and cell density of yeast strains with different ploidy. Sci Rep 13(1):1513 doi:10.1038/s41598-023-28800-z

Gallone B, Steensels J, Prahl T, Soriaga L, Saels V, Herrera-Malaver B, Merlevede A, Roncoroni M, Voordeckers K, Miraglia L, Teiling C, Steffy B, Taylor M, Schwartz A, Richardson T, White C, Baele G, Maere S, Verstrepen KJ (2016) Domestication and Divergence of Saccharomyces cerevisiae Beer Yeasts. Cell 166(6):1397–1410.e16 doi:10.1016/j.cell.2016.08.020

Garrison E, Marth G (2012) Haplotype-based variant detection from short-read sequencing. arXiv preprint arXiv:12073907:9–9 doi:arXiv:1207.3907 [q-bio.GN]

Haase SB, Reed SI (2002) Improved flow cytometric analysis of the budding yeast cell cycle. Cell cycle (Georgetown, Tex) 1(2):132–6 doi:10150107 [pii]

Hallahan BF (2024) One Hundred Years of Progress and Pitfalls: Maximising Heterosis through Increasing Multi-Locus Nuclear Heterozygosity. Biology (Basel) 13(10) doi:10.3390/biology13100817

Hallin J, Martens K, Young AI, Zackrisson M, Salinas F, Parts L, Warringer J, Liti G (2016) Powerful decomposition of complex traits in a diploid model. Nat Commun 7:13311 doi:10.1038/ncomms13311

Herbst RH, Bar-Zvi D, Reikhav S, Soifer I, Breker M, Jona G, Shimoni E, Schuldiner M, Levy AA, Barkai N (2017) Heterosis as a consequence of regulatory incompatibility. BMC Biology 15(1):38–38 doi:10.1186/s12915-017-0373-7

Hirota S, Nakayama Y, Ekino K, Harashima S (2024) Highly genomic instability of super-polyploid strains of Saccharomyces cerevisiae. J Biosci Bioeng 137(2):77–84 doi:10.1016/j.jbiosc.2023.11.009

Hochholdinger F, Yu P (2025) Molecular concepts to explain heterosis in crops. Trends Plant Sci 30(1):95–104 doi:10.1016/j.tplants.2024.07.018

Huxley C, Green ED, Dunbam I (1990) Rapid assessment of S. cerevisiae mating type by PCR. Trends in Genetics 6:236–236 doi:10.1016/0168-9525(90)90190-H

Kassambara A (2023) rstatix: Pipe-Friendly Framework for Basic Statistical Tests.

Kassambara A (2025) ggpubr: ‘ggplot2’ Based Publication Ready Plots.

Kolde R (2015) pheatmap : Pretty Heatmaps. R package version 108

Krogerus K, Arvas M, De Chiara M, Magalhães F, Mattinen L, Oja M, Vidgren V, Yue JX, Liti G, Gibson B (2016) Ploidy influences the functional attributes of de novo lager yeast hybrids. Applied Microbiology and Biotechnology 100(16):7203–7222 doi:10.1007/s00253-016-7588-3

Krogerus K, Fletcher E, Rettberg N, Gibson B, Preiss R (2021a) Efficient breeding of industrial brewing yeast strains using CRISPR/Cas9-aided mating-type switching. Applied Microbiology and Biotechnology doi:10.1007/s00253-021-11626-y

Krogerus K, Magalhães F, Castillo S, Peddinti G, Vidgren V, De Chiara M, Yue J-X, Liti G, Gibson B (2021b) Lager Yeast Design Through Meiotic Segregation of a Saccharomyces cerevisiae × Saccharomyces eubayanus Hybrid. Frontiers in Fungal Biology 2 doi:10.3389/ffunb.2021.733655

Krogerus K, Magalhães F, Vidgren V, Gibson B (2015) New lager yeast strains generated by interspecific hybridization. Journal of industrial microbiology & biotechnology 42(5):769–78 doi:10.1007/s10295-015-1597-6

Krogerus K, Magalhães F, Vidgren V, Gibson B (2017) Novel brewing yeast hybrids: creation and application. Applied Microbiology and Biotechnology 101(1):65–78 doi:10.1007/s00253-016-8007-5

Li H (2011) A statistical framework for SNP calling, mutation discovery, association mapping and population genetical parameter estimation from sequencing data. Bioinformatics 27(21):2987–2993 doi:10.1093/bioinformatics/btr509

Li H, Durbin R (2009) Fast and accurate short read alignment with Burrows-Wheeler transform. Bioinformatics 25(14):1754–1760 doi:10.1093/bioinformatics/btp324

Liu L, Jin M, Huang M, Zhu Y, Yuan W, Kang Y, Kong M, Ali S, Jia Z, Xu Z, Xiao W, Cao L (2021) Engineered Polyploid Yeast Strains Enable Efficient Xylose Utilization and Ethanol Production in Corn Hydrolysates. Front Bioeng Biotechnol 9:655272 doi:10.3389/fbioe.2021.655272

Loegler V, Friedrich A, Schacherer J (2024) Overview of the Saccharomyces cerevisiae population structure through the lens of 3,034 genomes. G3 (Bethesda) 14(12) doi:10.1093/g3journal/jkae245

Long Y, Han X, Meng X, Xu P, Tao F (2024) A robust yeast chassis: comprehensive characterization of a fast-growing Saccharomyces cerevisiae. mBio 15(2):e0319623 doi:10.1128/mbio.03196-23

Marchesan AN, Leal Silva JF, Maciel Filho R, Wolf Maciel MR (2021) Techno-Economic Analysis of Alternative Designs for Low-pH Lactic Acid Production. ACS Sustainable Chemistry & Engineering 9(36):12120–12131 doi:10.1021/acssuschemeng.1c03447

Marsit S, Henault M, Charron G, Fijarczyk A, Landry CR (2021) The neutral rate of whole-genome duplication varies among yeast species and their hybrids. Nat Commun 12(1):3126 doi:10.1038/s41467-021-23231-8

Merlini L, Dudin O, Martin SG (2013) Mate and fuse: how yeast cells do it. Open Biology 3(3):130008–130008 doi:10.1098/rsob.130008

Mertens S, Steensels J, Saels V, De Rouck G, Aerts G, Verstrepen KJ (2015) A large set of newly created interspecific Saccharomyces hybrids increases aromatic diversity in lager beers. Applied and environmental microbiology 81(23):8202–14 doi:10.1128/AEM.02464-15

Miller M, Zhang C, Chen ZJ (2012) Ploidy and Hybridity Effects on Growth Vigor and Gene Expression in Arabidopsis thaliana Hybrids and Their Parents. G3 (Bethesda) 2(4):505–13 doi:10.1534/g3.112.002162

Nikulin J, Krogerus K, Gibson B (2018) Alternative Saccharomyces interspecies hybrid combinations and their potential for low-temperature wort fermentation. Yeast 35(1):113–127 doi:10.1002/yea.3246

O’Donnell S, Yue J-X, Saada OA, Agier N, Caradec C, Cokelaer T, De Chiara M, Delmas S, Dutreux F, Fournier T, Friedrich A, Kornobis E, Li J, Miao Z, Tattini L, Schacherer J, Liti G, Fischer G (2023) Telomere-to-telomere assemblies of 142 strains characterize the genome structural landscape in Saccharomyces cerevisiae. Nature Genetics 55(8):1390–1399 doi:10.1038/s41588-023-01459-y

Peris D, Alexander WG, Fisher KJ, Moriarty RV, Basuino MG, Ubbelohde EJ, Wrobel RL, Hittinger CT (2020) Synthetic hybrids of six yeast species. Nature Communications 11(1):2085–2085 doi:10.1038/s41467-020-15559-4

Peris D, Moriarty RV, Alexander WG, Baker E, Sylvester K, Sardi M, Langdon QK, Libkind D, Wang Q-M, Bai F-Y, Leducq J-B, Charron G, Landry CR, Sampaio JP, Gonçalves P, Hyma KE, Fay JC, Sato TK, Hittinger CT (2017) Hybridization and adaptive evolution of diverse Saccharomyces species for cellulosic biofuel production. Biotechnology for Biofuels 10(1):78–78 doi:10.1186/s13068-017-0763-7

Peter J, De Chiara M, Friedrich A, Yue J-X, Pflieger D, Bergström A, Sigwalt A, Barre B, Freel K, Llored A, Cruaud C, Labadie K, Aury J-M, Istace B, Lebrigand K, Barbry P, Engelen S, Lemainque A, Wincker P, Liti G, Schacherer J (2018) Genome evolution across 1,011 Saccharomyces cerevisiae isolates. Nature 556(7701):339–344 doi:10.1038/s41586-018-0030-5

Plech M, de Visser JAGM, Korona R (2014) Heterosis Is Prevalent Among Domesticated but not Wild Strains of *Saccharomyces cerevisiae*. G3&#58; Genes|Genomes|Genetics 4(2):315–323 doi:10.1534/g3.113.009381

Preiss R, Fletcher E, Garshol LM, Foster B, Ozsahin E, Lubberts M, van der Merwe G, Krogerus K (2024) European farmhouse brewing yeasts form a distinct genetic group. Appl Microbiol Biotechnol 108(1):430 doi:10.1007/s00253-024-13267-3

Purcell S, Neale B, Todd-Brown K, Thomas L, Ferreira MAR, Bender D, Maller J, Sklar P, de Bakker PIW, Daly MJ, Sham PC (2007) PLINK: A Tool Set for Whole-Genome Association and Population-Based Linkage Analyses. The American Journal of Human Genetics 81(3):559–575 doi:10.1086/519795

Selmecki AM, Maruvka YE, Richmond Pa, Guillet M, Shoresh N, Sorenson AL, De S, Kishony R, Michor F, Dowell R, Pellman D (2015) Polyploidy can drive rapid adaptation in yeast. Nature 519(7543):349–352 doi:10.1038/nature14187

Shapira R, David L (2016) Genes with a Combination of Over-Dominant and Epistatic Effects Underlie Heterosis in Growth of Saccharomyces cerevisiae at High Temperature. Frontiers in Genetics 7 doi:10.3389/fgene.2016.00072

Shapira R, Levy T, Shaked S, Fridman E, David L (2014) Extensive heterosis in growth of yeast hybrids is explained by a combination of genetic models. Heredity 113(4):316–326 doi:10.1038/hdy.2014.33

Snoek T, Picca Nicolino M, Van den Bremt S, Mertens S, Saels V, Verplaetse A, Steensels J, Verstrepen KJ (2015) Large-scale robot-assisted genome shuffling yields industrial Saccharomyces cerevisiae yeasts with increased ethanol tolerance. Biotechnology for biofuels 8(1):32–32 doi:10.1186/s13068-015-0216-0

Steensels J, Meersman E, Snoek T, Saels V, Verstrepen KJ (2014a) Large-scale selection and breeding to generate industrial yeasts with superior aroma production. Applied and Environmental Microbiology 80(22):6965–6975 doi:10.1128/AEM.02235-14

Steensels J, Snoek T, Meersman E, Nicolino MP, Voordeckers K, Verstrepen KJ (2014b) Improving industrial yeast strains: Exploiting natural and artificial diversity. FEMS Microbiology Reviews 38(5):947–995 doi:10.1111/1574-6976.12073

Swinnen S, Goovaerts A, Schaerlaekens K, Dumortier F, Verdyck P, Souvereyns K, Van Zeebroeck G, Foulquie-Moreno MR, Thevelein JM (2015) Auxotrophic Mutations Reduce Tolerance of Saccharomyces cerevisiae to Very High Levels of Ethanol Stress. Eukaryot Cell 14(9):884–97 doi:10.1128/EC.00053-15

Takagi A, Harashima S, Oshima Y (1983) Construction and Characterization of Isogenic Series of Saccharomyces cerevisiae Polyploid Strains. Applied and Environmental Microbiology 45(3):1034–1040 doi:10.1128/aem.45.3.1034-1040.1983

Tarasov A, Vilella AJ, Cuppen E, Nijman IJ, Prins P (2015) Sambamba: fast processing of NGS alignment formats. Bioinformatics 31(12):2032–2034 doi:10.1093/bioinformatics/btv098

Tirosh I, Reikhav S, Levy AA, Barkai N (2009) A yeast hybrid provides insight into the evolution of gene expression regulation. Science 324(5927):659–662 doi:324/5927/659 [pii]10.1126/science.1169766

Tripp JD, Lilley JL, Wood WN, Lewis LK (2013) Enhancement of plasmid DNA transformation efficiencies in early stationary-phase yeast cell cultures. Yeast 30(5):191–200 doi:10.1002/yea.2951

Vittorelli N, Gomez-Munoz C, Andriushchenko I, Ollivier L, Agier N, Delmas S, Corbeau Y, Achaz G, Cosentino Lagomarsino M, Liti G, Llorente B, Fischer G (2025) Repeated losses of self-fertility shaped heterozygosity and polyploidy in yeast evolution. bioRxiv doi:10.1101/2025.09.12.675800

Winge O, Laustsen O (1938) Artificial species hybridization in yeast. CR Trav Lab Carlsberg, Ser Physiol 22:235–244

Wirth NT, Funk J, Donati S, Nikel PI (2023) QurvE: user-friendly software for the analysis of biological growth and fluorescence data. Nat Protoc 18(8):2401–2403 doi:10.1038/s41596-023-00850-7

Xie Z-X, Mitchell LA, Liu H-M, Li B-Z, Liu D, Agmon N, Wu Y, Li X, Zhou X, Li B, Xiao W-H, Ding M-Z, Wang Y, Yuan Y-J, Boeke JD (2018) Rapid and Efficient CRISPR/Cas9-Based Mating-Type Switching of Saccharomyces cerevisiae. G3 Genes|Genomes|Genetics 8(1):173–183 doi:10.1534/g3.117.300347

Zavaleta V, Perez-Traves L, Saona LA, Villarroel CA, Querol A, Cubillos FA (2024) Understanding brewing trait inheritance in de novo Lager yeast hybrids. mSystems 9(12):e0076224 doi:10.1128/msystems.00762-24

Zhu Y, Yogiswara S, Willekens A, Gerardin A, Lavigne R, Goossens A, Pinheiro VB, Dai Z, Verstrepen KJ (2025) Beyond CEN.PK -parallel engineering of selected S. cerevisiae strains reveals that superior chassis strains require different engineering approaches for limonene production. Metab Eng 91:276–289 doi:10.1016/j.ymben.2025.04.011

Zörgö E, Chwialkowska K, Gjuvsland AB, Garré E, Sunnerhagen P, Liti G, Blomberg A, Omholt SW, Warringer J (2013) Ancient Evolutionary Trade-Offs between Yeast Ploidy States. PLoS Genetics 9(3):e1003388–e1003388 doi:10.1371/journal.pgen.1003388

