## Supplementary Figures and Tables for "Influence of ploidy and genetic background on stress tolerance of intraspecific yeast hybrids": Supplementary_Figures.pdf

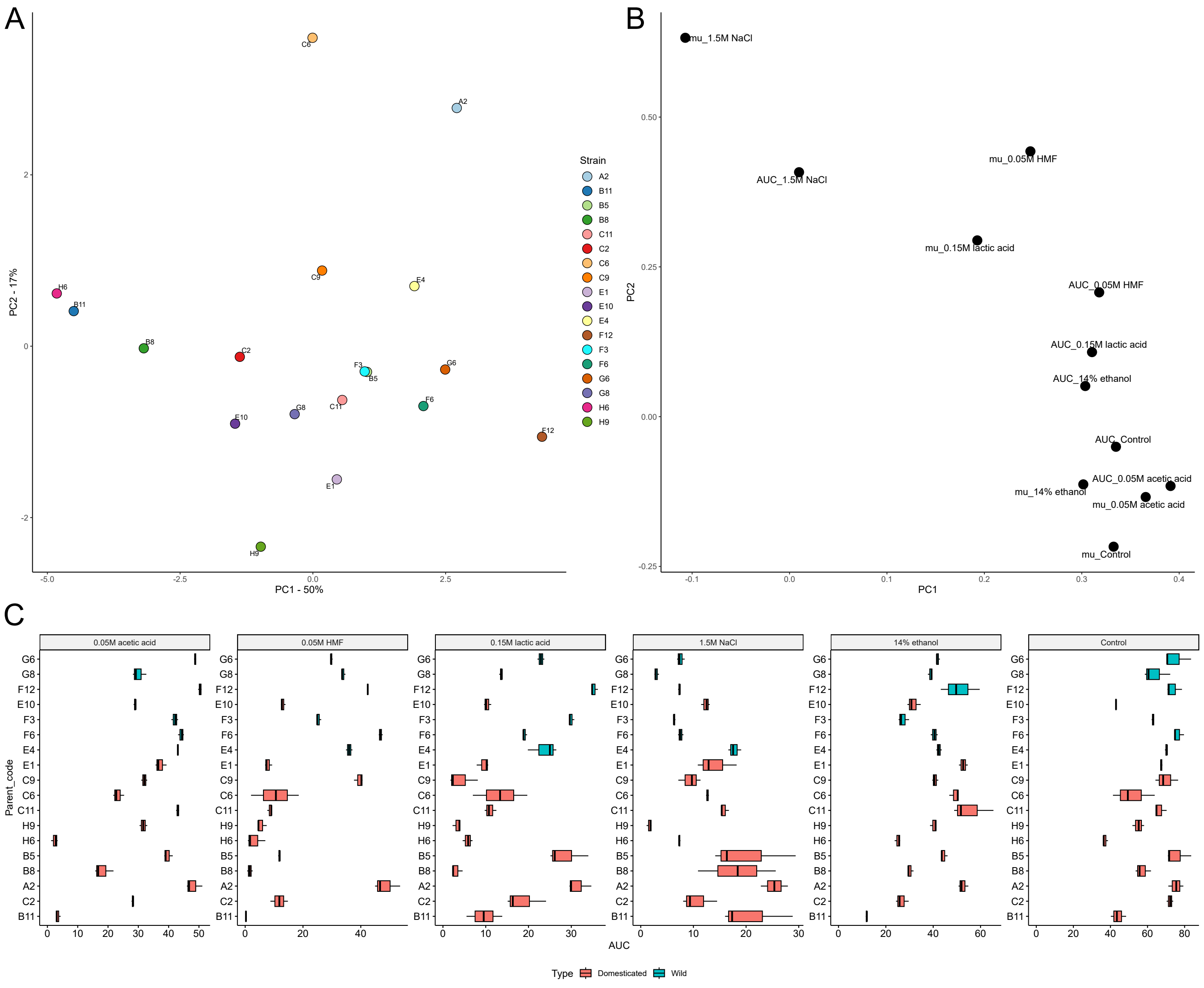


Supplementary Figure S1. Growth of the wild-type parent strains in different media. (A-B) Principal component analysis of the growth data (AUC and μ) in the six different growth media, with the (A) scores plot to the left, and the (B) loadings plot to the right. The first two principal components explain 67% of the variance. (C) Boxplots of the area under curve (AUC) in the wild-type parent strains in the six different growth media. Strains are ordered based on the phylogenetic relationship between the parents (maximum likelihood phylogenetic tree based on SNPs at 105677 sites, rooted with strain G8 as outgroup). Boxes are colored cyan if the parent strain is considered wild, red if considered domesticated.

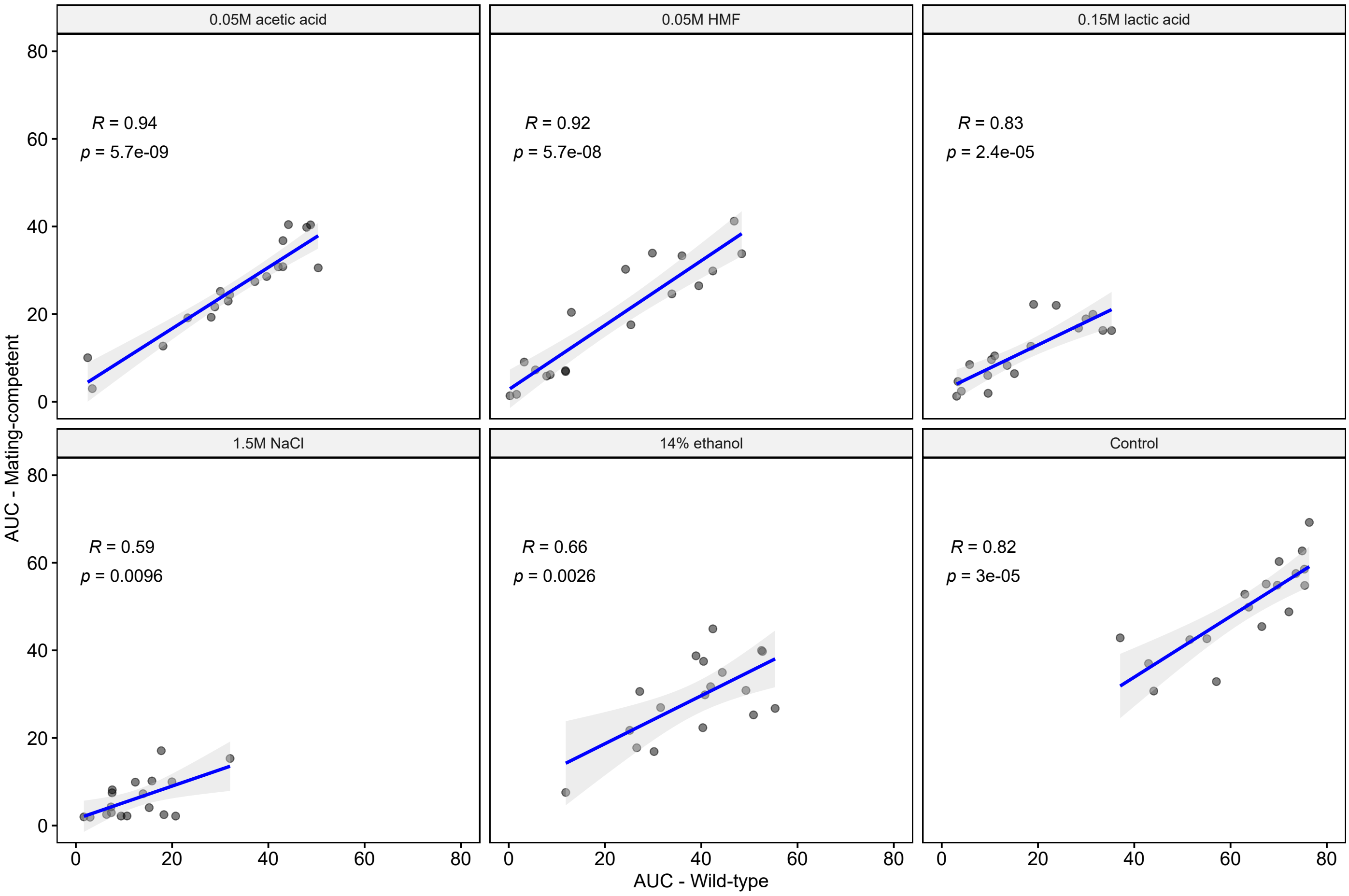


Supplementary Figure S2. The correlation between AUC in the wild-type and mating-competent parent strains in the different growth media. *R* is the Pearson correlation coefficient.

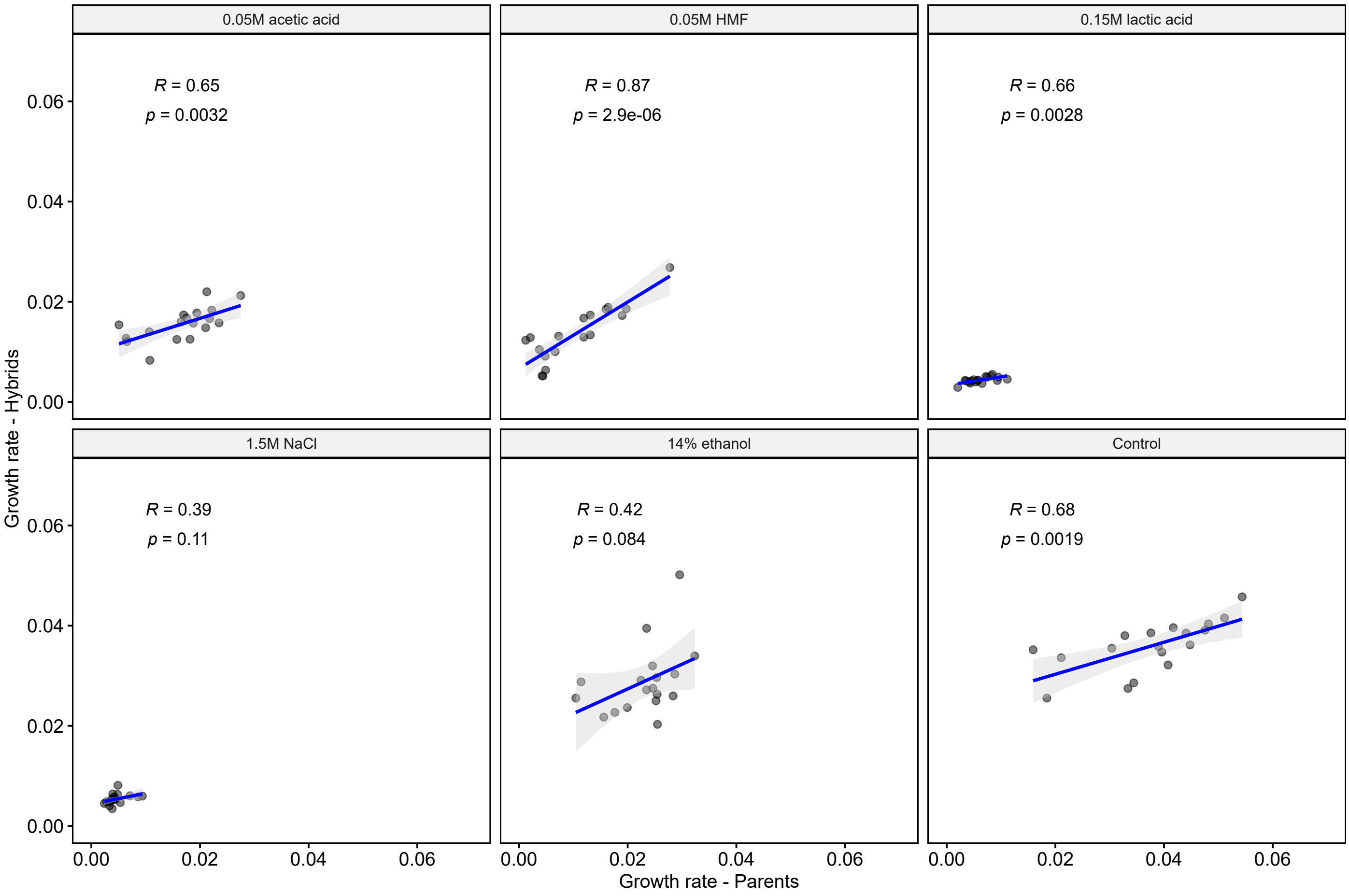


Supplementary Figure S3. The correlation between μ (growth rate) in the parent strains and hybrids grouped by parent strain in the different growth media. *R* is the Pearson correlation coefficient.

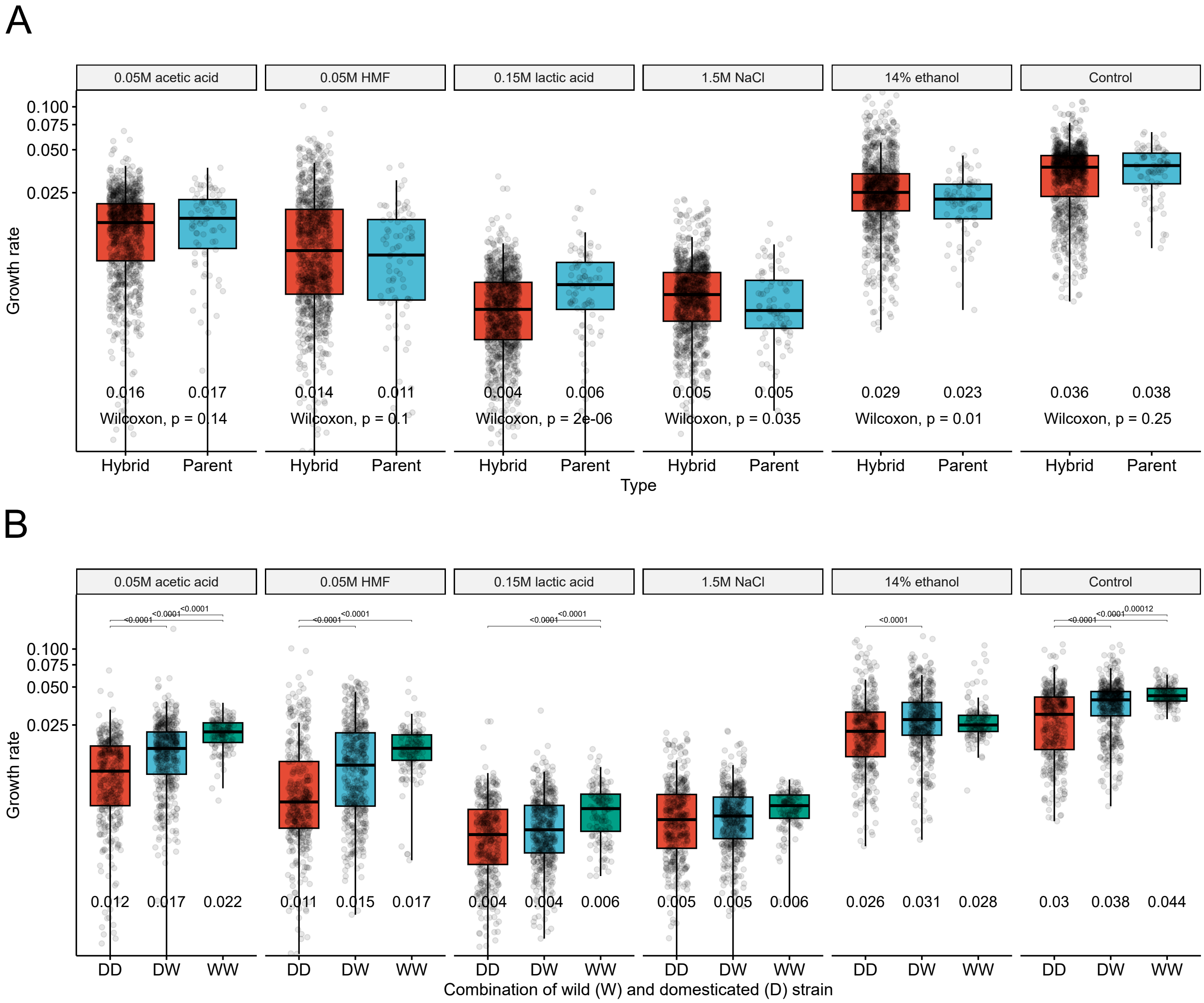


Supplementary Figure S4. Growth comparison of the hybrid and parent strains in different media. (A) Boxplots of the growth rate (μ) in the hybrid (red) and parent (blue) strains in the six different growth media. Mean values of each group are displayed under the boxes. Statistical difference between the hybrid and parent strains in each media was tested by an unpaired Wilcoxon test. (B) Boxplots of the growth rate (μ) in the hybrid strains as grouped by parent strain combination in the six different growth media. DD (red): both parent strains were domesticated, DW (blue): one parent strain was domesticated, and one was wild, WW (green): both parent strains were wild. Statistical difference between the groups in each media was tested by one-way ANOVA and Tukey's post-hoc test, and *p*-values are shown above the boxes.

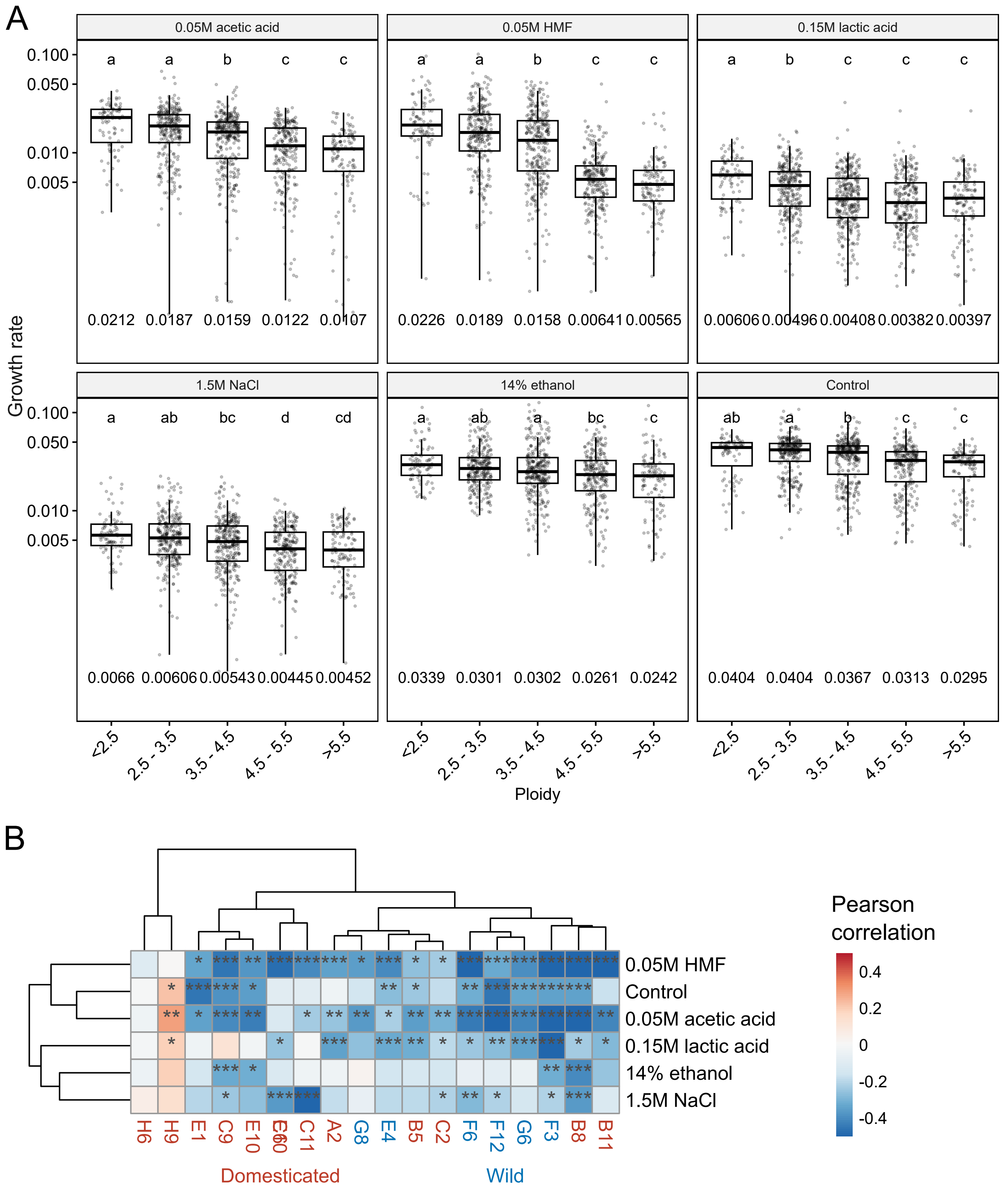


Supplementary Figure S5. Influence of ploidy on the growth of the hybrids in different media. (A) Boxplots of the growth rate (μ) in the hybrid strains as grouped by ploidy in the six different growth media. The mean value of each group is shown below the boxes. Different letters above the boxes within each growth media indicate significant differences ($p < 0.05$) as determined by one-way ANOVA and Tukey's post-hoc test. (B) A heatmap visualizing the Pearson correlation coefficient between ploidy and μ in the hybrids as grouped by parent strain and growth media. Red and blue colors indicate a positive and negative correlation coefficient, respectively. The *p*-values from multiple comparisons were corrected using the Benjamini-Hochberg procedure. Asterisks indicate the *p*-values as follows: * $p < 0.05$, ** $p < 0.01$, and *** $p < 0.001$.

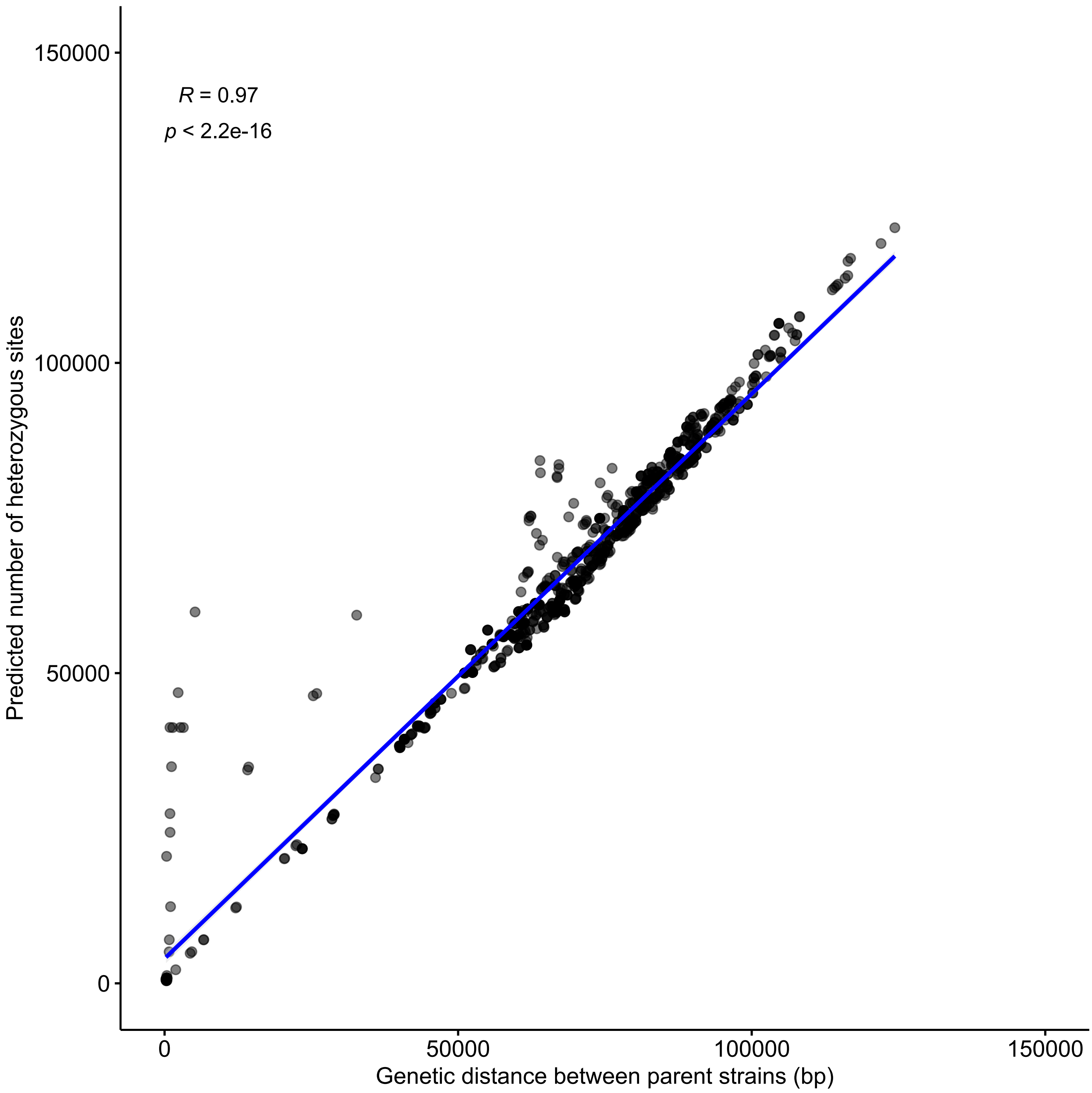


Supplementary Figure S6. The correlation between the genetic distance between parent strains (bp) and the estimated amount of heterozygous sites in the hybrids. Genetic distance was estimated based on the sum of IBS0 and IBS1 alleles in each pairwise cross as estimated in PLINK based on SNPs detected in the mating-competent parent strains of each hybrid. *R* is the Pearson correlation coefficient. Hybrids with low genetic distance between parent strains, yet high predicted heterozygosity, are inbred strains where the parent strains were heterozygous.

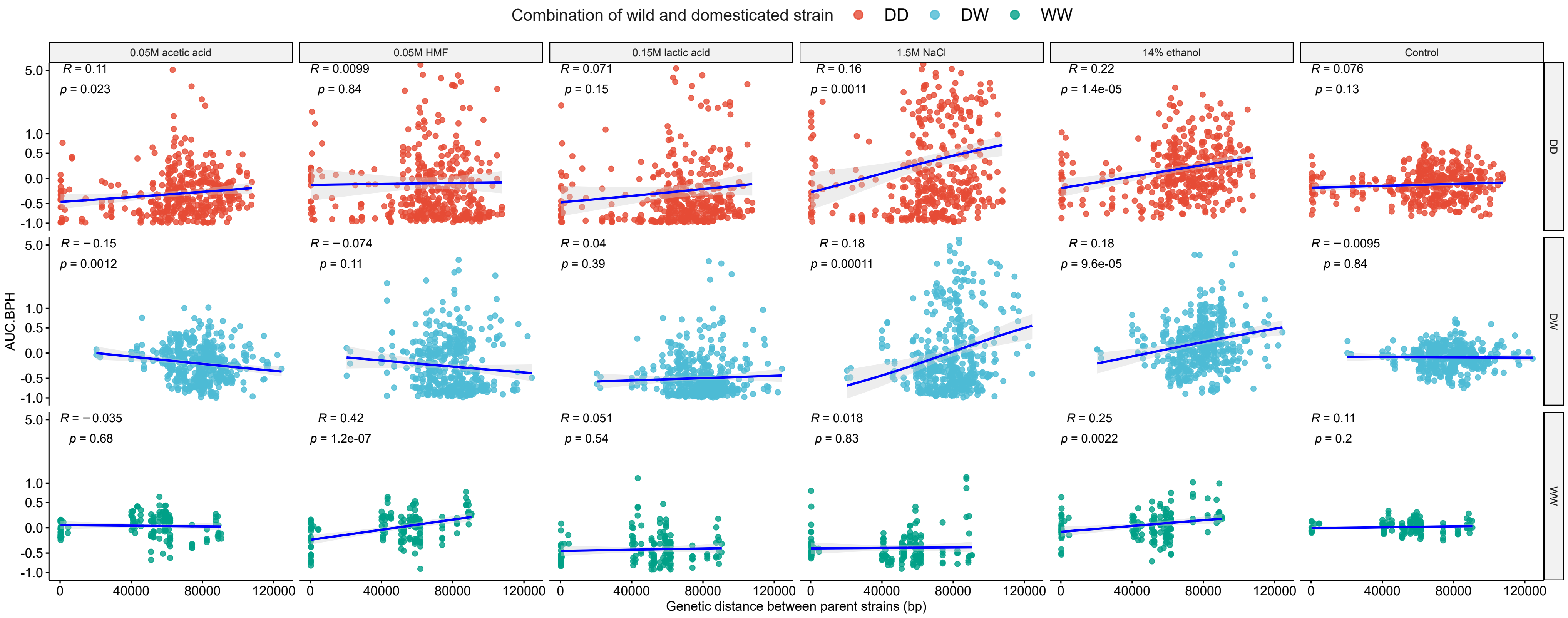


Supplementary Figure S7. Influence of genetic distance between parent strains (bp) on the best-parent heterosis based on area under curve (AUC.BPH) in the hybrid strains as grouped by parent strain combination in the six different growth media. DD (red): both parent strains were domesticated, DW (blue): one parent strain was domesticated, and one was wild, WW (green): both parent strains were wild. Genetic distance was estimated based on the sum of IBS0 and IBS1 alleles in each pairwise cross as estimated in PLINK based on SNPs detected in the mating-competent parent strains of each hybrid. *R* is the Pearson correlation coefficient.
